# A hierarchical transcriptional network controls appressorium-mediated plant infection by the rice blast fungus *Magnaporthe oryzae*

**DOI:** 10.1101/2020.02.05.936203

**Authors:** Míriam Osés-Ruiz, Magdalena Martin-Urdiroz, Darren M. Soanes, Michael J. Kershaw, Neftaly Cruz-Mireles, Guadalupe Valdovinos-Ponce, Camilla Molinari, George R. Littlejohn, Paul Derbyshire, Frank L. H. Menke, Barbara Valent, Nicholas J. Talbot

## Abstract

Rice blast is a pervasive and devastating disease that threatens rice production across the world. In spite of its importance to global food security, however, the underlying biology of plant infection by the blast fungus *Magnaporthe oryzae* remains poorly understood. In particular, it is unclear how the fungus elaborates a specialised infection cell, the appressorium, in response to surface signals from the rice leaf. Here, we report the identification of a network of temporally co-regulated transcription factors that act downstream of the Pmk1 mitogen-activated protein kinase pathway to regulate gene expression during appressorium-mediated plant infection. We have functionally characterised this network of transcription factors and demonstrated the operation of a hierarchical transcriptional control system. We show that this tiered regulatory mechanism involves Pmk1-dependent phosphorylation of the Hox7 homeobox transcription factor, which represses hyphal-associated gene expression and simultaneously induces major physiological changes required for appressorium development, including cell cycle arrest, autophagic cell death, turgor generation and melanin biosynthesis. Mst12 then regulates gene functions involved in septin-dependent cytoskeletal re-organisation, polarised exocytosis and effector gene expression necessary for plant tissue invasion.

---

Rice blast disease, caused by the fungus *Magnaporthe oryzae*, is one of the most important threats to global food security ^1^. The disease starts when spores of the fungus, called conidia, land on the hydrophobic surface of a rice leaf. Conidia attach strongly to the leaf cuticle and germinate rapidly to produce a short germ tube, that soon differentiates into a dome-shaped, infection cell called an appressorium ^1–3^. The appressorium develops enormous turgor of up to 8.0MPa, as a result of glycerol accumulation in the cell which draws water into the appressorium by osmosis, thereby generating hydrostatic pressure ^4^. Glycerol is maintained in the appressorium by a thick layer of melanin in the cell wall, which reduces its porosity ^4,5^. Development of the appressorium is tightly linked to cell cycle control and autophagic re-cycling of the contents of the conidium ^6–8^. Appressorium turgor is monitored by a sensor kinase, Sln1, and once a threshold of pressure has been reached ^9^, septin GTPases are recruited to the appressorium pore, where they form a toroidal, hetero-oligomeric complex that scaffolds cortical F-actin at the base of the appressorium ^10^, leading to protrusive force generation to enable a penetration peg to pierce the cuticle of the rice leaf and gain entry to host tissue. Once inside the leaf, invasive hyphae colonize the first epidermal cell before seeking out pit field sites, where plasmodesmata are situated, through which the fungus invades neighbouring plant cells ^11^. *M. oryzae* actively suppresses plant immunity using a battery of fungal effector proteins delivered into plant cells ^12^. After five days disease lesions appear, from which the fungus sporulates to colonize neighbouring plants.

Formation of an appressorium by *M. oryzae* requires a conserved <u>p</u>athogenicity <u>m</u>itogen-activated protein <u>k</u>inase, called Pmk1 ^13^. Pmk1 mutants are unable to form appressoria and cause plant infection, even when plants have been previously wounded ^13^. Instead spores of *Δpmk1* mutants produce undifferentiated germlings that undergo several rounds of mitosis and septation ^13,14^. Pmk1 is responsible for lipid and glycogen mobilization to the appressorium and the onset of autophagy in the conidium ^4,8,15,16^, but also plays a role in cell-to-cell movement by *M. oryzae* during invasive growth. Chemical genetic inactivation of Pmk1, prevents the fungus from moving through pit fields between rice cells ^11^. Pmk1 is therefore a global regulator of both appressorium development and fungal invasive growth. Little, however, is known regarding the means by which this kinase exerts such a significant role in the developmental biology of *M. oryzae*. So far, for example, only one transcriptional regulator, Mst12, has been identified which appears to act downstream of Pmk1. Mst12 mutants form appressoria normally, but these are non-functional and *Δmst12* mutants are unable to cause rice blast disease ^17^.

In this study we set out to identify the mechanism by which major transcriptional changes are regulated during appressorium development by *M. oryzae*. During a time-course of appressorium development on an inductive hydrophobic surface, we identified major temporal changes in gene expression that occur in response to surface hydrophobicity and which require the Pmk1 MAPK, or Mst12 transcription factor. In this way we were able to identify cellular pathways associated with appressorium morphogenesis and maturation, including a group of Pmk1-regulated transcription factor-encoding genes. We functionally characterised this group of putative regulators and were able to define a hierarchy of genetic control. We show that the Hox7 homeodomain transcription factor is phosphorylated by Pmk1 and co-ordinates regulation of genes associated with development of a functional appressorium, while simultaneously repressing hyphal growth, and then interacting with Mst12, which regulates a second distinct population of genes necessary for re-polarisation of the appressorium and invasive growth.

## Results

### The Pmk1 MAPK is necessary for appressorium development in response to surface hydrophobicity

Appressorium development by *M. oryzae* can be readily induced under laboratory conditions by incubating spores on hard hydrophobic surfaces. When conidia are incubated on a hydrophobic surface they form appressoria within 6h and septins accumulate at the base of the incipient appressorium, forming a ring structure by 8h ^10^, as shown in Fig. 1a. At the same time, a single round of mitosis occurs, and the conidium undergoes autophagic cell death and degradation of the three conidial nuclei ^8^, so that a single nucleus is present in the mature appressorium by 14h (Fig. 1a-e), By contrast, when *M. oryzae* conidia are incubated on a hydrophilic surface, a completely different morphogenetic program occurs in which long germlings develop that do not differentiate into appressoria and conidia do not undergo autophagy (Fig. 1a, b) ^6^. Multiple rounds of mitosis occur and by 24h ~50% of germlings contain greater than 4 nuclei (Fig. 1b). Appressorium development in *M. oryzae* requires the Pmk1 signalling pathway ^13,16^, as shown in Fig 1c. Pmk1 mutants do not develop appressoria and instead, form undifferentiated germlings unable to cause plant infection (Fig. 1d). Strikingly, the germlings produced by *Δpmk1* mutants undergo multiple rounds of mitosis and conidia do not undergo autophagic cell death (Fig. 1e), as previously reported ^6^. Therefore, the response of the fungus to a hydrophilic surface closely mirrors that of mutants lacking the Pmk1 MAPK. This is consistent with the role of Pmk1 as a regulator of appressorium development in response to an inductive, hydrophobic surface.

**Figure 1.**
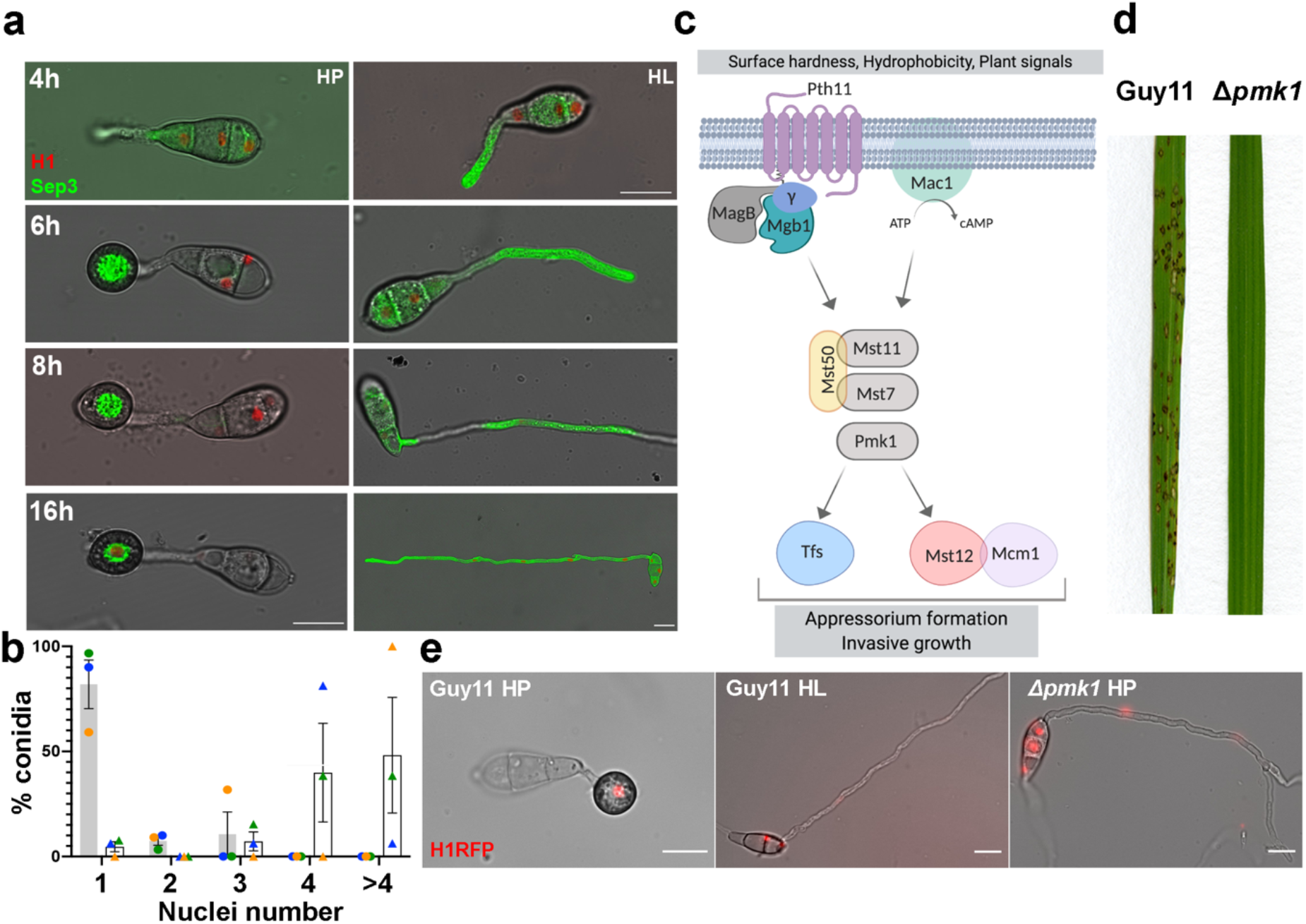
The Pmk1 MAP kinase signalling pathway regulates appressorium development in response to surface hydrophobicity: **a.** Micrographs to show development of a wild type *M. oryzae* strain Guy11 expressing histone H1-RFP nuclear marker and septin Sep3-GFP, inoculated on a hydrophobic (HP) or hydrophilic (HL) surface. Scale Bar = 10 μm. **b**. Bar chart to show proportion of Guy11 germlings containing, 1, 2, 3, 4 or more nuclei following incubation for 24h on HP or HL surfaces. **c.** Schematic representation of the Pmk1 MAPK signalling pathway in *M. oryzae*. Dissociation of Mgb1 causes activation of Mst11 (MAPKKK), which activates Mst7 (MAPKK) and, ultimately, Pmk1 (MAPK). The MAPK signalling complex is scaffolded by Mst50. Mst7 activates Pmk1 which regulates a series of uncharacterised transcription factors, as well as Mst12 ^56,57^ **d.** Rice blast disease symptoms of Guy11 and the Δ*pmkl* mutant. Rice seedlings of cultivar CO-39 were spray-inoculated with conidial suspensions of equal concentrations of each *M. oryzae* strain and grown for 5 days. **e.** Live cell imaging to show nuclear number and the presence/absence of conidial autophagic cell death of Guy11 on HP, Δ*pmkl* mutant on HP and Guy11 on HL surfaces. Each strain expressed the H1-RFP nuclear marker.

### The global transcriptional response of *M. oryzae* to surface hydrophobicity

We decided to determine the global transcriptional response of *M. oryzae* in response to surface hydrophobicity and compare this to transcriptional changes associated with loss of Pmk1. To do this, we isolated total RNA from a wild type strain of *M. oryzae*, Guy11, during a time course of appressorium development on hydrophobic glass coverslips (hereafter referred to as the HP surface) and on hydrophilic Gelbond membranes (hereafter, called the HL surface), from 0h-24h after inoculation. In parallel, we incubated conidia of a Δ*pmk1* mutant on the HP surface in an identical time course, as shown in Fig. 2a. We then carried out RNA sequencing analysis (RNA-seq) and identified *M. oryzae* genes differentially expressed in response to surface hydrophobicity, or the presence/absence of Pmk1. For this, we used a p-value adjusted for false discovery rate (padj < 0.01), as shown in Fig2 b and c. We observed 3917 differentially expressed *M. oryzae* genes in response to surface hydrophobicity (HP vs HL) and 6333 genes differentially expressed in a Δ*pmk1* mutant compared to the isogenic wild type, Guy11 (Supplementary Tables 1-2). By comparing these gene sets we identified 3555 genes differentially expressed in response to both a HP surface and the presence of Pmk1, a small set of 362 Pmk1-independent genes differentially expressed in response to the HP surface, and 2778 Pmk1-dependent genes that did not show differential expression between HP and HL surfaces, as shown in Fig 2d (Supplementary Table 3). We conclude that 90% of genes that are differentially expressed by *M. oryzae* in response to surface hydrophobicity are also Pmk1-dependent.

**Figure 2.**
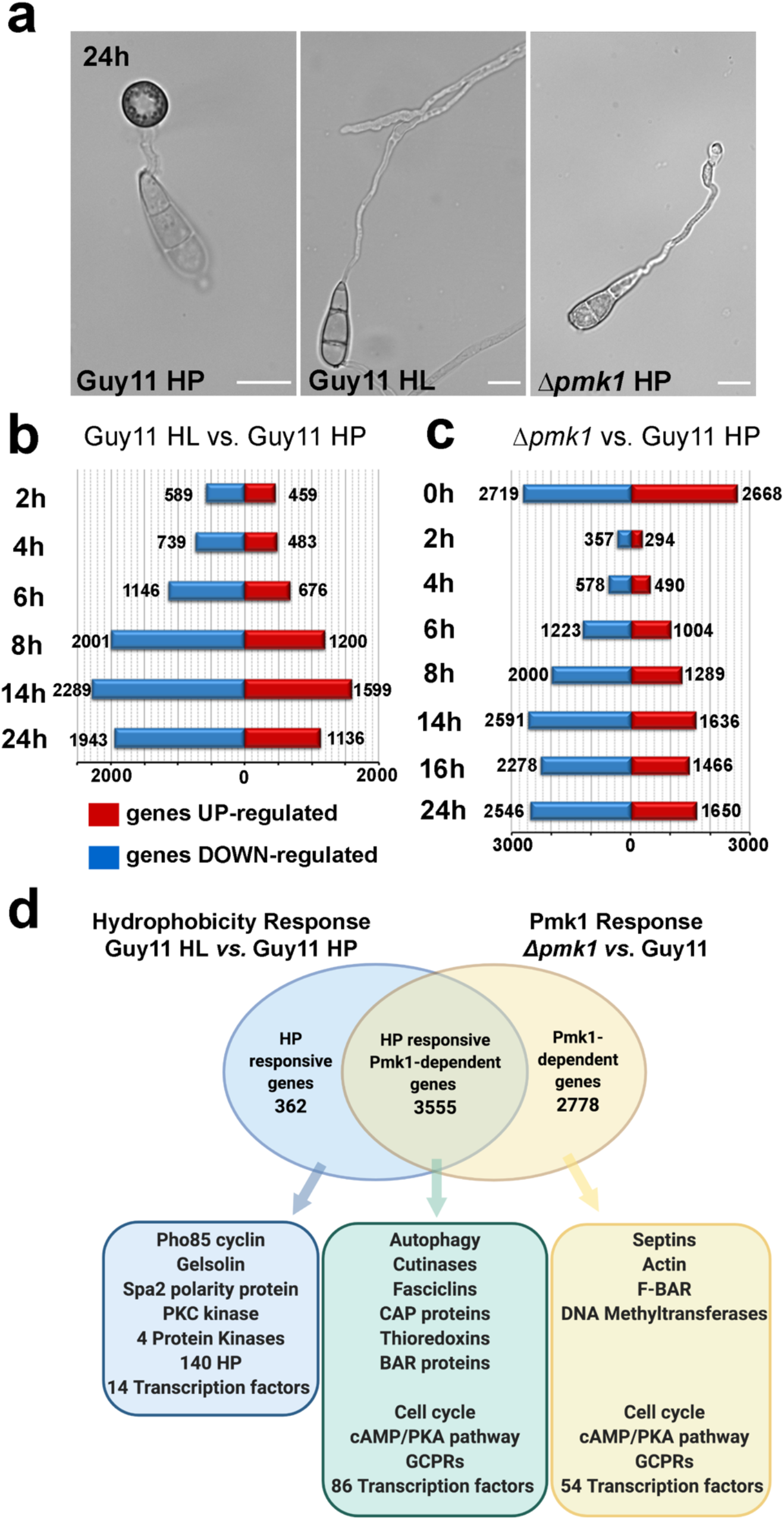
Global comparative transcriptional profile analysis to define the response of *M. oryzae* to surface hydrophobicity and the presence/absence of the Pmk1 MAPK. **a.** Bright field microscopy to show germ tube extension and appressorium formation of Guy11 on HP surface, Guy11 on HL surface, and a Δ*pmk1* mutant on HP surface. Scale bar = 10 μm. **b**. Bar charts to show number of up-regulated genes (red) (padj< 0.01, mod_lfc >1) and down-regulated genes (blue) (padj< 0.01, mod_lfc < −1) of Guy11 in response to incubation on HP or HL surfaces between 2h and 24h. **c**. Bar charts to show number of up-regulated genes (red) (padj< 0.01, mod_lfc >1) and down-regulated genes (blue) (padj< 0.01, mod_lfc < −1) in a Δ*pmk1* mutant compared Guy11 following incubation on HP surface from 0h - 24h. **d.** Venn diagram illustrating overlapping sets of genes showing at least 2-fold differential expression in at least two time points (padj<0.01, mod_lfc>1 or mod_lfc<-1) in Guy11 in response to incubation on HP or HL surfaces, or between the Δ*pmk1* mutant compared to Guy11 on HP surface. Three distinct populations of genes could be identified, as HP-surface responsive (blue), HP-response and Pmk1-dependent (green), or Pmk1-dependent only (yellow).

We analysed the set of 3555 HP/Pmk1-dependent genes to define functions associated with appressorium morphogenesis. Differential regulation of genes required for a number of physiological processes previously implicated in appressorium morphogenesis was observed, including autophagy– responsible for conidial cell death and appressorium function ^6,8^ –which is clearly dependent on Pmk1 and is a response to HP surfaces. Autophagy-associated genes are highly expressed during early stages of appressorium development and down-regulated from 8 h onwards (Supplementary Figs. 1 and 2). Similarly, cell cycle-related genes show differential regulation in response to HP surfaces and many are Pmk1-dependent (Supplementary Figure 2). Genes encoding cyclins; the main cyclin dependent kinase *CDC28*; its positive regulator *MIH1*, and negative regulator *SWE1* ^18^ all showed expression peaks at 4h-6h, coincident with appressorium morphogenesis (Supplementary Fig. 3-4). By contrast, cyclin-associated gene expression was delayed and CDK-related gene expression oscillated abnormally in *Δpmk1* mutants. We also found that genes required for the DNA damage response (DDR) pathway ^19^ were mis-regulated in *Δpmk1* mutants. *DUN1* gene expression, for example, was altered during appressorium development. We additionally found evidence that the significant repertoire of <u>G</u>-<u>p</u>rotein <u>c</u>oupled <u>r</u>eceptors (GPCRs) in *M. oryzae* ^20^ are involved in appressorium morphogenesis. A total of 50 of the 79 known GPCR-encoding genes in *M. oryzae* (63%) were differentially regulated in response to the HP surface and 48 were Pmk1-dependent (Supplementary Fig. 5). Analysis of differentially expressed genes implicated further physiological processes associated with appressorium morphogenesis, including 14 acetyltransferases, 13 ABC transporters, 2 BAR domain proteins, 95 Major facilitator superfamily transporter, 24 protein kinases and 86 transcription factors, 3 fasciclins and 6 cutinases in this group, as shown in Supplementary Table 1.

We next analysed 2778 Pmk1-dependent, HP surface-independent genes. Interestingly, there was considerable overlap in predicted gene functions (see Supplementary Table 2), but we observed more changes associated with appressorium maturation, such as cytoskeletal re-modelling, with extensive differential regulation of genes encoding actin-binding proteins, cofilin, coronin, the F-actin capping protein, Fes/CIP4, and EFC/F-BAR domain proteins (Supplementary Fig. 6). Differential expression of septins and their associated regulators was also observed (Supplementary Fig 5 b, c, d), consistent with the requirement for septin-dependent, F-actin re-modelling in appressorium function ^9,10^.

We conclude that Pmk1 acts as a global regulator of gene expression in *M. oryzae* in response to surface hydrophobicity, with more than 90% of HP-responsive genes requiring Pmk1. However, the role of the MAPK also extends to appressorium maturation, following initial development of the infection cell.

### Defining the role of the Pmk1-dependent transcriptional regulator Mst12 in control of appressorium gene expression

Having revealed that Pmk1 is required for expression of more than 6000 genes during appressorium development, we decided to investigate the role of known downstream regulators, with the aim of establishing a hierarchy of genetic control during plant infection by *M. oryzae*. Previously, the Ste12-like-zinc finger transcription factor, Mst12 was identified as being regulated by Pmk1 ^17^. Δ*mst12* mutants still form appressoria, but are unable to cause plant infection ^17^, as shown in Fig 3a-b and Supplementary Fig. 5a. To further define the likely function of Mst12, we carried out live cell imaging of appressoria of a Δ*mst12* mutant and found the mutant impaired in its ability to undergo septin-dependent, F-actin re-modelling at the appressorium pore (Fig. 3c), required for plant infection ^10^. We observed disorganisation of the microtubule network in the appressorium in the Δ*mst12* mutant (Fig. 3c) and, consistent with impairment in septin organisation, mis-localisation of the PAK-related kinase, Chm1, the Ezrin/ Radixin/Moesin, Tea1, the Bim-amphiphysin-Rvs (BAR) domain-containing protein, Rvs167, and the Staurosporine and Temperature sensitive-4, Stt4 lipid kinase (Fig. 3d) ^9,10^. We conclude that Mst12 is necessary for septin-dependent re-polarisation of the appressorium.

**Figure 3.**
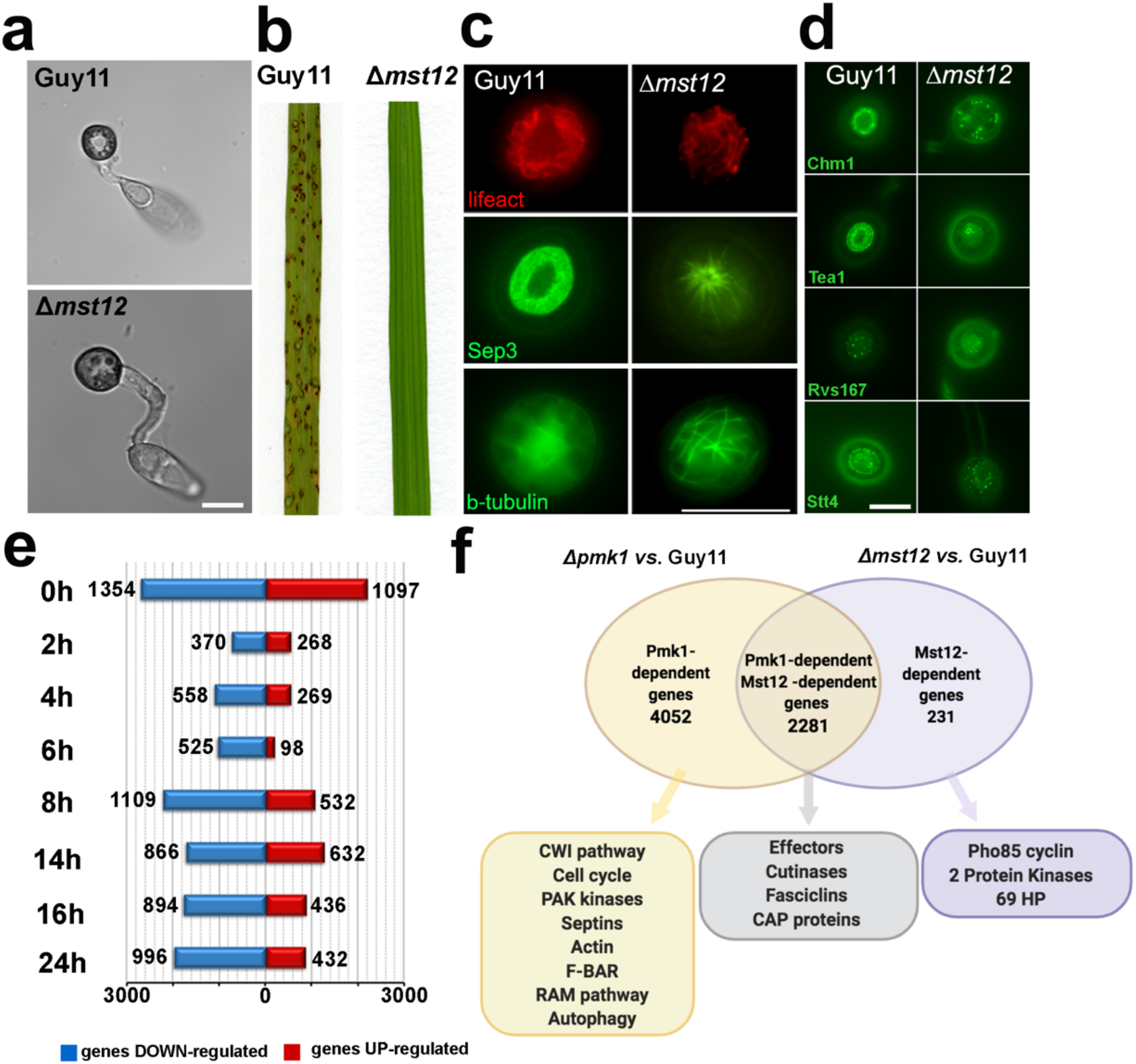
Functional analysis and comparative global transcriptional profile analysis in response to the presence/absence of *M. oryzae* Mst12 transcription factor. **a.** Micrograph to show appressorium development of Guy11 and a Δ*mst12* mutant after incubation for 24h on HP surface. Scale Bar = 10 μm. **b.** Rice blast disease symptoms of Guy11 and a Δ*mst12* mutant. Rice seedlings of cultivar CO-39 were spray-inoculated with conidial suspensions of equal concentrations of each *M. oryzae* strain and incubated for 5 days. **c.** Live cell imaging to show cellular localization of Lifeact-RFP, Sep3-GFP and β-tubulin-GFP in appressorium pore of Guy11 and Δ*mst12*, following incubation on HP surface for 24 h. Scale Bar = 10 μm. **d.** Micrographs to show cellular localization of Chm1-GFP, Tea1-GFP, Rvs167-GFP and Stt4-GFP in appressorium pore of Guy11 and Δ*mst12* following incubation on HP surface for 24 h. Scale Bar = 10 μm. **e.** Bar chart to show number of up-regulated genes (red) (padj< 0.01, mod_lfc >1) and down-regulated genes (blue) (padj< 0.01, mod_lfc < −1) in a Δ*mst12* mutant compared to Guy11 following incubation on a HP surface between 0h and 24h. **e.** Venn diagram illustrating overlapping sets of genes showing at least 2-fold differential expression in at least two time points (padj<0.01, mod_lfc>1 or mod_lfc<-1) between a Δ*pmk1* mutant compared to Guy11, incubated on HP surface and Δ*mst12* mutant compared to Guy11 incubated on HP surface. Three distinct populations of genes could be identified, as Pmk1-dependent (yellow), Pmk1 and Mst12-dependent (grey), or Mst12-dependent only (purple).

As a consequence of the clear role for Mst12 in regulating the latter stages of appressorium maturation, we performed RNA-seq analysis during a time course of appressorium development by the *Δmst12* mutant and then compared its global pattern of gene expression to those genes regulated by Pmk1 (Supplementary Tables 4-5). In this way, we reasoned that we would be able to define both early-acting Pmk1-dependent genes and a later group of gene functions requiring both Pmk1 and Mst12. We identified 2512 differentially expressed genes in the *Δmst12* mutant with significant changes in gene expression occurring during conidial germination and particularly after 8h, during the onset of appressorium maturation (Fig. 3e). When this Mst12-regulated gene set was compared to the wider Pmk1-requiring gene set, we identified 4052 genes that are Pmk1-dependent, but Mst12-independent, associated with initial stages of appressorium development, while 2281 maturation-associated genes are regulated by both Pmk1 and Mst12 (Fig. 3f). Only 231 Mst12-dependent genes were independent of Pmk1 control (Fig. 3f). A clear example of a Pmk1-dependent, Mst12-independent gene family were the melanin biosynthetic genes, required for appressorium function ^21^, which showed maximum gene expression at 6h in Guy11, were not expressed at all in *Δpmk1* mutants, but were expressed in *Δmst12* mutants at 8h, when appressoria form in the mutant (Supplementary Fig. 7).

To investigate genes associated with appressorium maturation, we identified gene functions regulated by both Pmk1 and Mst12, from among the 2281 differentially expressed genes. The cutinase gene family, previously implicated in appressorium morphogenesis and plant infection ^22^, for example, was dependent on both Pmk1 and Mst12 and a family of fasciclins– membrane-associated glycoproteins involved in cell adhesion ^23,24^ –were also regulated by both Pmk1 and Mst12 (Supplementary Fig. 8). A total of 436 genes encoding putatively secreted proteins were also part of the Pmk1-Mst12-dependent group, including 7 known effector genes (Supplementary Fig. 9). Two of these effectors, Bas2 and Bas3, have been reported to be Pmk1-dependent during invasive growth of the fungus in rice tissue ^11^. To provide direct evidence of Pmk1-dependent regulation of effector gene expression, we expressed Bas2p:GFP in a *pmk1^as^* mutant (Supplementary Fig. 9e). This mutant allows conditional inactivation of the Pmk1 MAPK in the presence of the ATP analogue drug Na-PP1 ^11^. When Na-PP1 was applied to the *pmk1^as^* mutant expressing Bas2pGFP, we observed loss of Bas2-GFP fluorescence, suggesting that activity of Pmk1 is necessary for expression of the effector during appressorium maturation, prior to plant infection.

Given the significant effect of Mst12 on expression of secreted proteins, such as effectors, we reasoned that the Δ*mst12* mutant might be impaired in secretory functions. Septin recruitment to the base of the appressorium is, for example, necessary for organisation of the octameric exocyst complex, required for polarised exocytosis ^25^, so we expressed a GFP fusion protein of the exocyst component Sec6 in both *Δmst12* and Guy11 (Supplementary Fig. 8). In wild type appressoria, Sec6-GFP formed a ring structure, while in *Δmst12* mutants it was completely absent (Supplementary Fig. 8). We also localised the Flp2 fasciclin in Guy11 and the *Δmst12* mutant. Flp2-GFP localises to the plasma membrane during appressorium development at 8h and by 24h to the base of the appressorium in a Mst12-dependent manner (Supplementary Fig. 8). We conclude that Pmk1 and Mst12 are both necessary for expression of a large set of genes associated with appressorium maturation, including genes associated with polarised exocytosis and effector genes which serve roles in plant tissue invasion.

### Identifying the hierarchy of transcriptional regulation required for appressorium development by *M. oryzae*

Our results suggested that Pmk1 acts as a global regulator of appressorium morphogenesis, while Mst12 regulates a specific sub-set of genes associated with cytoskeletal reorganisation, re-polarisation and effector secretion. This strongly suggested that other transcription factors must act downstream of Pmk1. We therefore determined the total number of putative transcription factor-encoding genes differentially regulated at least 2-fold (padj <0.01, mod_lfc >1 or mod_lfc <-1) in at least two time points in *Δpmk1* and/or *Δmst12* mutants, compared to Guy11. We found 140 such transcription factor genes, of which 95 were associated with appressorium morphogenesis and dependent on Pmk1, and 45 associated with appressorium maturation and dependent on both Pmk1 and Mst12 (Fig. 4a). We plotted heatmaps for each gene set (Fig. 4b, 4c and 4d) which enabled us to identify a specific clade of transcription factor-encoding genes severely down-regulated in the Δ*pmk1* mutant, which we called Clade 4 (Fig. 4c). Clade 4 consists of 15 putative transcription factors (Supplementary Tables 6) including nine Zn_2_Cys_6_ transcription factors, some of which have been implicated in stress responses (Fzc64, Fzc52, Fzc41 and Fzc30), conidial germination (Fzc50) or appressorium formation (Fzc75)^26^. Three transcription factors were, however, uncharacterized and we named them <u>R</u>elated to <u>P</u>mk1 <u>P</u>athway (RPPs) genes; *RPP1* (MGG_10212), *RPP2* (MGG_09276) and *RPP3* (MGG_07218). We also found two Zn_2_Cys_6_ and fungal specific domain transcription factor-encoding genes that we called *RPP4* (MGG_07368) and *RPP5* (MGG_8917), as well as the *PIG1* transcription factor-encoding gene (MGG_07215), previously identified as a regulator of melanin biosynthesis in *M. oryzae* ^27^. *ALCR* (MGG_02129), a homologue of *AlcR* from *Aspergillus nidulans* responsible for activation of the ethanol-utilization pathway through activation of *AlcA* and *AldA* ^28^ and a homeobox domain transcription factor-encoding gene *HOX7* (MGG_12865), previously shown to be involved in appressorium formation and pathogenicity ^29^, were also part of Clade 4. These transcription factor genes all showed strong down-regulation in a *Δpmk1* mutant from 4h onwards (Fig. 4e), the time when appressoria first develop, and were altered in expression in the *Δmst12* mutant at later time points, consistent with the delay in appressorium formation observed in the mutant (Fig. 4f).

**Figure 4:**
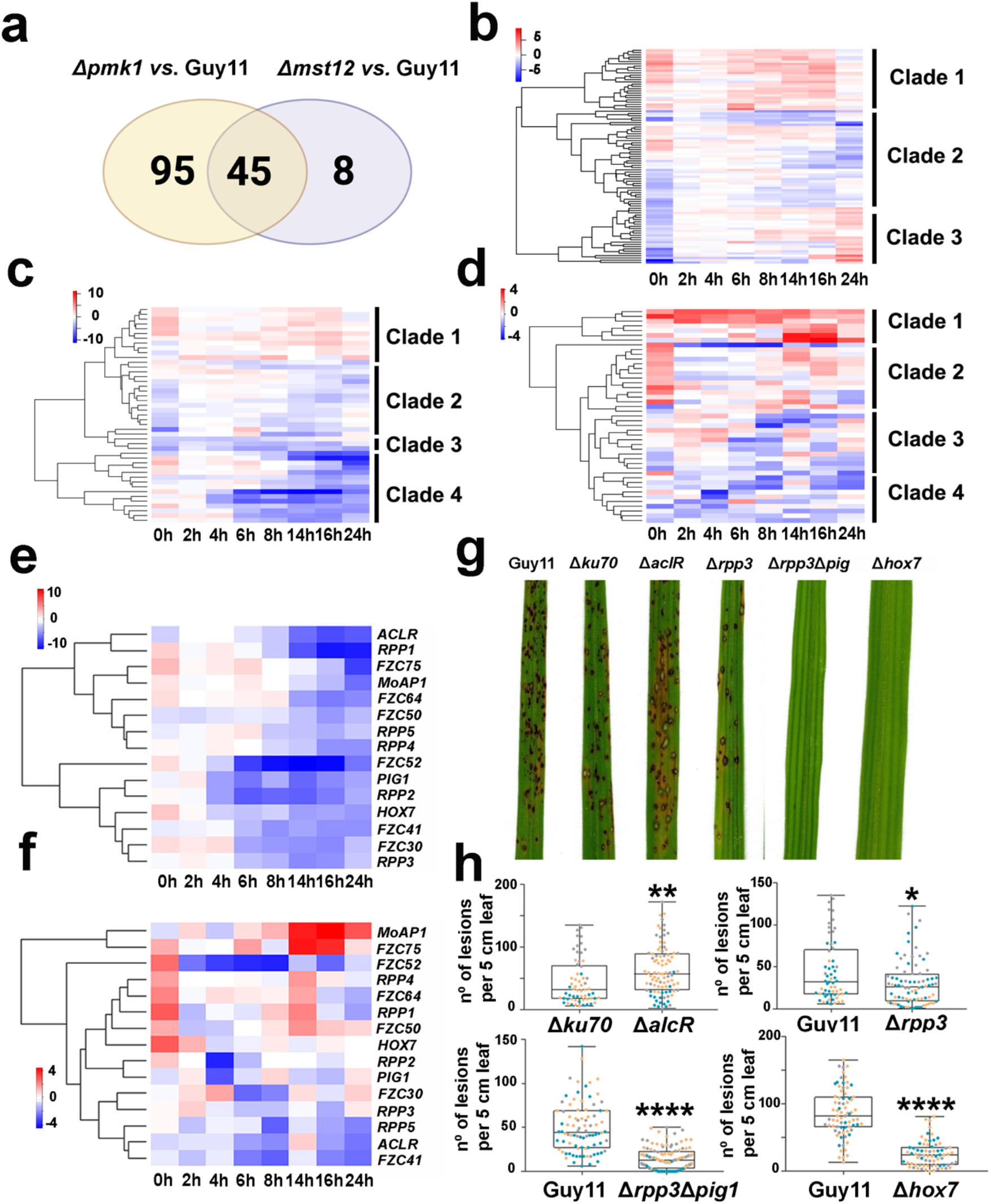
Defining the hierarchy of transcriptional control during appressorium development by *M. oryzae*. **a.** Venn diagram showing overlapping sets of *M. oryzae* putative transcription factor-encoding genes showing at least 2-fold differential expression in at least two time points (padj<0.01, mod_lfc>1 or mod_lfc<-1) between Δ*pmk1* and Guy11 (yellow), Δ*mst12* and Guy11 (purple), or both Δ*pmk1*, Δ*mst12* and Guy11 (grey). **b.** Heatmap showing temporal pattern of relative transcript abundance of 95 transcription factor-encoding genes between a Δ*pmk1* mutant and Guy11. **c.** Heatmap showing temporal pattern of relative transcript abundance of 45 transcription factor-encoding genes between a Δ*pmk1* mutant and Guy11. **d.** Heatmap showing temporal pattern of relative transcript abundance of 53 transcription factor-encoding genes between a Δ*mst12* mutant and Guy11. **e.** Heatmap showing temporal patterns of transcript abundance of Clade 4 transcription factor-encoding genes differentially regulated in a Δ*pmk1* mutant compared to Guy11. **f.** Heatmap showing temporal patterns of transcript abundance of Clade 4 transcription factor-encoding genes differentially regulated in a Δ*mst12* mutant compared to Guy11 (blue= down-regulated in the mutant; red= up-regulated in the mutant). **g.** Rice blast disease symptoms of Δ*aclR, Δrpp3*, Δ*rpp3Δpig1*, and Δ*hox7* mutants compared to Guy11 and a Δ*ku70* mutant. Rice seedlings of cultivar CO-39 were spray-inoculated with conidial suspensions of equal concentrations of each *M. oryzae* strain incubated for 5 days. **h.** Box and whisker plots to show number of rice blast disease lesions per 5 cm leaf in pathogenicity assays. Data points are shown in whisker plots which show 25^th^/75^th^ percentiles, the median and the minimum and maximum values by the ends of the whiskers. A two-tailed nonparametric Mann Whitney statistical test was conducted. Statistical analysis showed: Δ*aclR vs. Δku70* p=0.0017, n= 3 biological replicates; Δ*rpp3 vs*. Guy11 p=0.0131, n= 4 biological replicates; Δ*rpp3Δpig1 vs*. Guy11 p<0.0001, n= 4 biological replicates; Δ*hox7 vs*. Guy11 p<0.0001, n= 4 biological replicates.

To test whether these clade 4-associated transcription factor-encoding genes play important roles in appressorium development and plant infection we carried out targeted gene deletions to generate isogenic *ΔaclR, Δrpp1, Δrpp2, Δrpp4, Δrpp5, Δrpp3, Δrpp3Δpig1 and Δhox7* null mutants in either *Δku70* (a mutant of Guy11 lacking the non-homologous DNA end-joining pathway, that facilitates efficient homologous recombination to create null mutants ^6^ or Guy11 (Supplementary Fig. 10). We decided to generate a double mutant for *RPP3* and *PIG1* genes because both genes are regulators of melanin biosynthesis with likely overlapping functions (Supplementary Fig. 10). We inoculated 21-day old seedlings of the blast-susceptible rice cultivar CO-39 with conidial suspensions of each mutant and quantified disease symptoms after 5 days (Fig. 4g-h; Supplementary Fig. 11). Guy11 and *Dku70* were able to cause plant infection as expected but by contrast, the Δ*hox7* mutant was non-pathogenic (Fig. 4g), while both the *Δrpp3* and Δ*rpp3Δpig1* mutants were significantly reduced in their ability to cause disease. Conversely, the *ΔaclR* mutant showed a slight increase in the number of rice blast lesions compared to the wild type (Fig. 4g-h).

### The Hox7 homeobox transcription factor is directly regulated by the Pmk1 MAP kinase

To see if the loss of rice blast symptoms in Clade 4 transcription factor mutants was due to impairment in appressorium development, we tested if mutant strains could develop appressoria after 24h on HP surfaces (Supplementary Fig. 11). All the mutants developed appressoria indistinguishable from Guy11, except *Δhox7* mutants which developed immature non-melanised appressoria that re-germinated to produce hypha-like structures (Fig. 5a-b). Because loss of Hox7 affected appressorium development, we hypothesised that the transcription factor might act in a distinct manner compared to Mst12 and the remaining transcription factors of Clade 4. Hox7 is part of a family of six homeobox-domain (PF00046) transcription factor-encoding genes and two homeobox KN domain (PF05920) encoding-genes previously identified, but its relationship to the Pmk1 MAPK is unknown ^29^. To investigate the function of Hox7, we carried out RNA-seq analysis of a *Δhox7* mutant germinated on HP surfaces at 14 h and compared the global pattern of gene expression to that observed in *Δpmk1* and *Δmst12* mutants (Fig. 5c; Supplementary Tables 7-8). Hox7 controls expression of 4211 genes, of which 2332 are down-regulated (padj <0.01, mod_lfc<-1) and 1879 up-regulated (padj <0.01, mod_lfc > 1), as shown in Fig. 5c. To understand the hierarchy of genetic control between Pmk1, Mst12 and Hox7, we determined overlapping gene sets of differentially regulated genes between *Δhox7, Δpmk1*, and *Δmst12* mutants, compared to Guy11 (Fig. 5c). Pmk1 and Hox7 share 1942 differentially expressed genes, while Mst12 and Hox7 share only 169 differentially expressed genes (Fig. 5c). Our analysis revealed 709 genes differentially expressed in *Δpmk1, Δmst12* and *Δhox7* mutants, which when plotted in a heatmap revealed a very high level of similarity in the pattern of gene expression between *Δpmk1* and *Δhox7* mutants, in contrast to the transcriptional signature of the *Δmst12* mutant (Supplementary Fig. 12). Hox7 may therefore act directly downstream of Pmk1 to regulate a strongly overlapping set of genes. To test this idea, we first looked at expression of Clade 4 transcription factors (Table 3). When we plotted a heatmap showing moderated log fold changes in expression of these transcription factor genes, we confirmed that their pattern of gene expression was very similar in *Δpmk1* and *Δhox7* mutants (Fig. 5d). We therefore widened our analysis to the 1942 genes differentially regulated by both Pmk1 and Hox7, which revealed a strongly overlapping pattern of transcriptional regulation (Supplementary Fig. 12). Analysis of cellular processes regulated by both Pmk1 and Hox7 revealed the RAM pathway, melanin biosynthesis, autophagy and cell cycle control (Supplementary Fig. 12). To investigate how cell cycle control and autophagy are regulated by Hox7, we first looked at expression of cyclin genes Cln3, Clb2 and Clb3; CDK-related genes Mih1, Cdc28, Swe1 and Cks1; and DNA Damage Response (DDR) pathway-related genes Cds1, Dun1 and Chk1. Expression of all these genes was altered in the *Δhox7* mutant (Fig. 5e), as were autophagy-related genes (Fig. 5f). To investigate the potential role of Hox7 in cell cycle control and autophagic cell death, we therefore introduced a Histone H1-RFP nuclear marker into Guy11 and the *Δpmk1* mutant, and a H1-GFP marker into *Δmst12* and *Δhox7* mutants. We inoculated conidia on HP surfaces and monitored nuclear number over 24 h (Fig. 5g). The *Δhox7* mutant contained 4 or more nuclei by 24 h, resembling *Δpmk1*, whereas the *Δmst12* mutant resembled Guy11 with a single nucleus in the appressorium. Hox7 is therefore likely to be required for cell cycle arrest that occurs following mitosis in the germ tube. Furthermore, this experiment showed that conidia of both *Δhox7* and *Δpmk1* mutants did not collapse by 24h in the same way as a *Δmst12* mutant and the wild type Guy11. This is consistent with regulation of autophagy-related genes requiring both Pmk1 and Hox7 (Fig. 5f). When considered together, these findings provide evidence that Hox7 coordinates cell cycle progression and autophagy during appressorium development, probably acting as a negative regulator of cell cycle progression and an activator of autophagy-mediated conidial cell death, which are necessary to repress hyphal growth and progress appressorium differentiation.

**Figure 5:**
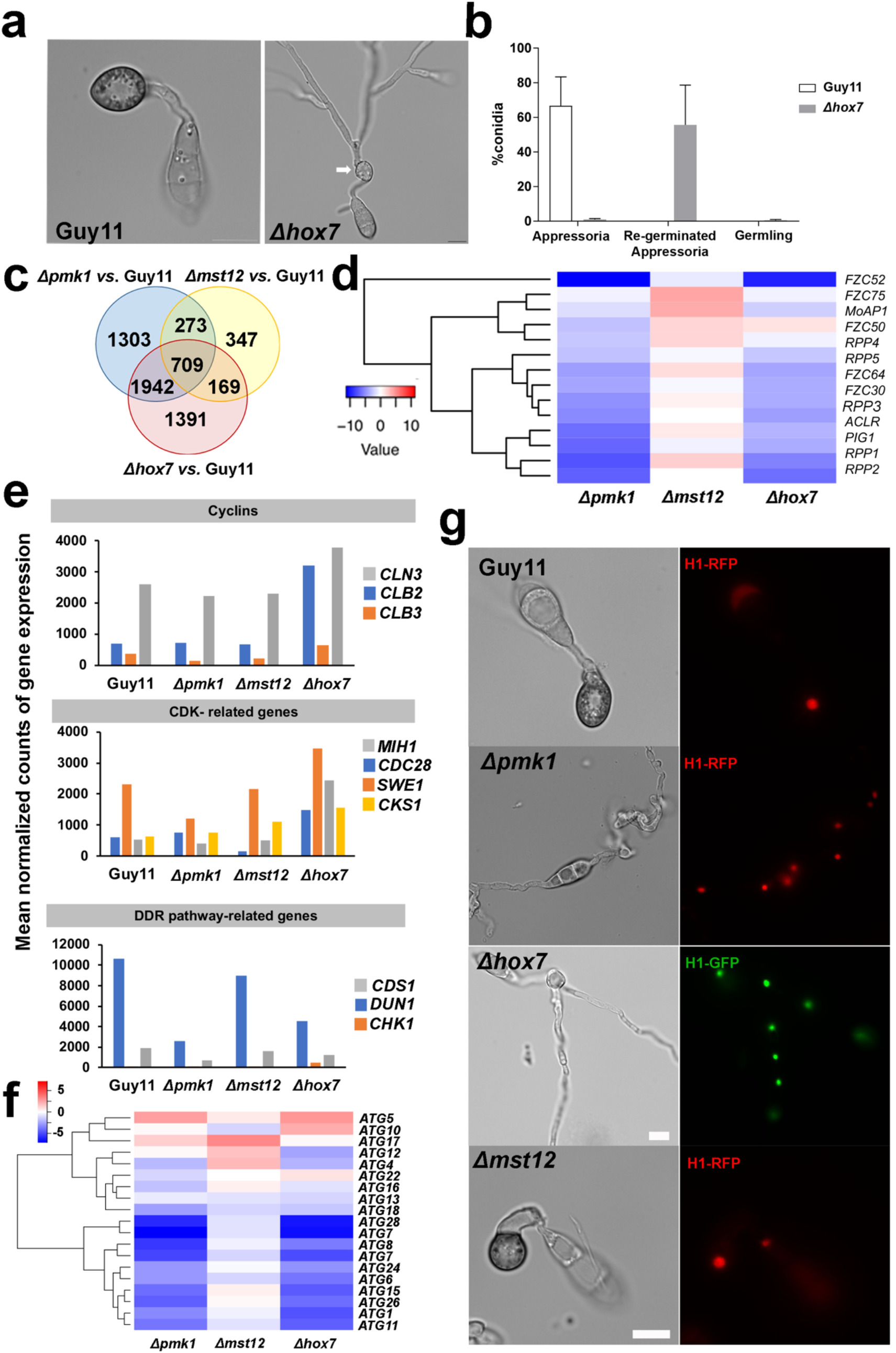
Characterisation of the Hox7 homeobox transcription factor and its role in the regulation of gene expression during appressorium development by *M. oryzae*. **a.** Bright field microscopy of appressorium development by Guy11 and a Δ*hox7* mutant following incubation on HP surface for 24h. Scale Bar = 10 μm. Arrow shows re-germination of an incipient appressorium and hyphal elongation. **b.** Bar chart to show defects in appressorium development of Δ*hox7* compared to Guy11 on HP surface at 24h (n=3 biological replicates; 50-100 conidia per replicate). **c.** Venn diagram to show overlapping sets of genes showing at least 2-fold difference (padj<0.01, mod_lfc>1 or mod_lfc<-1) between Guy11 and the Δ*pmk1* (blue), Δ*mst12* (yellow) and Δ*hox7* (red) mutants, respectively, during 14h timecourse of appressorium development. **d.** Heatmap to show relative transcript abundance of clade 4-associated transcription factors in Δ*pmk1*, Δ*mst12* and Δ*hox7* mutants compared to Guy11 **e.** Bar chart to show mean normalized counts of gene expression of cyclins, CDK-related genes and <u>D</u>NA <u>d</u>amage <u>r</u>esponse (DDR) pathway-related genes during appressorium development in Guy11, Δ*pmk1*, Δ*mst12* and Δ*hox7* mutants. **f.** Heatmap to show relative transcript abundance of autophagy-related genes in Δ*pmk1*, Δ*mst12* and Δ*hox7* mutants compared to Guy11. **g.** Live cell imaging to show nuclear dynamics in Guy11, Δ*pmk1, Δhox7* and Δ*mst12* mutants each expressing H1-GFP, after 24h germination on HP surface. Scale Bar = 10 μm.

### Hox7 is a direct target of the Pmk1 MAP kinase in *M. oryzae*

To define the function of the Pmk1 MAP kinase more precisely and, in particular, to understand its relationship with Hox7, we carried out discovery phosphoproteomics. We carried out appressorium development assays in *Δpmk1* and *Δhox7* mutants and Guy11 at 6h on HP surfaces using identical conditions to our RNA-seq analysis. Phosphoproteins were purified and analysed by tandem mass spectrometry. This revealed Pmk1-dependent phosphorylation of proteins associated with cell cycle control, including Dun1 and Far1, and autophagy-related proteins, including Atg13 and Atg26, and components of the cAMP-dependent protein kinase A pathway ^30,31^. Some of these processes are also dependent on the presence of Hox7 (Fig. 6a). We observed that Hox7 was phosphorylated at serine 158 (Fig. 6a-b) and therefore carried out parallel reaction monitoring (PRM), which revealed that Pmk1-dependent phosphorylation of Hox7 occurs 2h after conidial germination (Fig. 6c). We also tested whether Hox7 has the capacity to interact with Pmk1 and Mst12 *in vitro*. We constructed bait vectors for Pmk1 and Mst12 and prey vectors for Mst12 and Hox7 and tested their ability to interact in a yeast two hybrid experiment and found that Hox7 is able to interact weakly with Pmk1 and Mst12 (Fig. 6d). Our RNA-seq data furthermore showed that *HOX7* is highly expressed in Guy11 and *Δmst12* at 6h and 8h, respectively (Supplementary Fig. 12). Hox7 therefore functions at the point when an appressorium is formed, suggesting that Hox7 acts downstream of Pmk1, perhaps as a repressor of hyphal-like growth once the appressorium develops (Fig. 6e).

**Figure 6:**
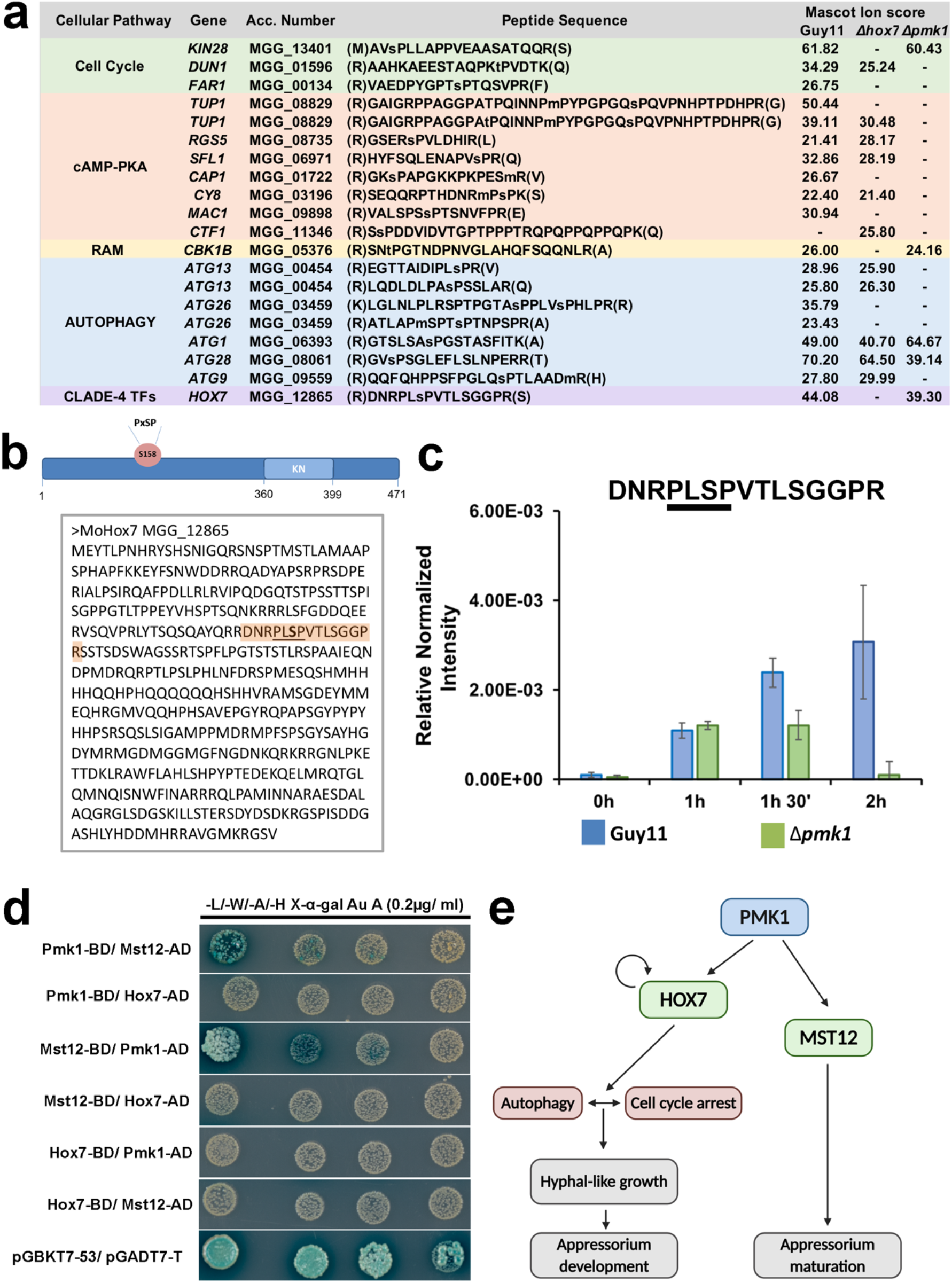
Phosphoproteomic analysis reveals Pmk1-dependent phosphorylation of Hox7 in *M. oryzae*. **a.** Classification of phosphoproteins identified by discovery phosphoproteomics in Guy11, Δ*pmk1* and Δ*hox7* mutants during appressorium development for 6h on HP surface. The specific peptide sequence containing proline-directed phosphorylation either at a serine or threonine residue for each accession number is shown with the corresponding mascot ion score. **b**. Predicted amino acid sequence of Hox7 protein to show serine 158 residue. Sequence of Hox7 protein to show identification of a MAP kinase phosphorylation motif with phosphorylation at serine 158. **c.** Parallel Reaction Monitoring (PRM) showing Pmk1-depemdent phosphorylation at serine 158 of Hox7 during appressorium development up to 2h after conidial germination. **d.** Yeast-two-hybrid assay to show interaction of Hox7 homeobox transcription factor with Pmk1 kinase and Mst12 transcription factor. Co-transformed Y2H Gold strains with bait (BD) and prey (AD) vectors in all possible combinations and positive control (pGBKT7-53 and pGADT7-T) were grown in double drop out media and quadruple dropout media supplemented with X- α-gal and Aureobasidin A. Images represent n= 3 biological replicates. **e.** Model to illustrate hierarchical control of gene expression by the Pmk1 MAPK and associated transcription factors Hox7 and Mst12. Hox7 is directly phosphorylated by Pmk1 and regulates autophagy and cell cycle arrest to promote appressorium development. Mst12 regulates appressorium maturation and re-polarisation.

## Discussion

Understanding how fungal pathogens are able to infect their host plants is critical if durable control strategies are to be developed to control crop diseases ^32^. The rice blast fungus has emerged as a useful model system to study plant infection processes because of its relative experimental tractability and, in particular, the generation of mutants unable to infect host plants ^33,34^. It has long been known, for example, that the Pmk1 MAPK signalling pathway is essential for controlling appressorium development ^13^ and many other determinants of plant infection have also been defined ^33–35^. Furthermore, transcriptional profile analysis has been used to define physiological processes involved in rice blast infections, such as control of autophagy, lipid metabolism, and melanin biosynthesis ^36^. However, what has remained completely unknown is how the global changes in gene expression essential for elaboration of an appressorium by *M. oryzae* are actually controlled.

In this study we have identified the global transcriptional signature associated with appressorium development by the rice blast fungus in response to surface hydrophobicity, and then used comparative transcriptome analysis using *Δpmk1* and *Δmst12* mutants, to define a set of Pmk1-dependent transcription factors involved in regulation of appressorium morphogenesis. In particular, we have demonstrated that Pmk1 regulates the Hox7 homeobox transcription factor to enable cell cycle arrest and autophagic conidial cell death, which are essential pre-requisites for appressorium-mediated infection ^8^. Furthermore, this has enabled us to determine the role of Mst12, which regulates septin-dependent F-actin re-modelling and re-polarisation of the appressorium, as well as controlling secretion of early acting effector proteins delivered into plant tissue at the point of plant infection. Collectively, this has enabled the formulation of the model presented in Fig. 6e.

Hox7 emerges from this study as a vital regulator of appressorium development. Hox7 was previously reported to be necessary for plant infection ^29^ but its function was unknown. We have now provided evidence that Hox7 is a direct target of Pmk1. Hox7, for example, contains a MAP kinase phosphorylation motif and is phosphorylated at serine 158 in a Pmk1-dependent manner (Fig. 5). Homeobox transcription factors regulate morphogenesis in many organisms, acting as both activators or repressors of genes during multicellular development ^37^. They were first described, for instance, as developmental-switch genes in the fruit fly *Drosophila melanogaster* and subsequently, in many other eukaryotes ^38 39^. Homeobox transcription factors have also been extensively studied in cancer biology, where they have been implicated in the control of autophagy and cell cycle regulation. In glioblastomas, for example, HoxC9 acts as a negative regulator of autophagy by controlling activation of beclin1 by regulating the transcription of the death associated protein kinase1 DAPK1 ^40^ and in neuroblastoma cells, HoxC9 interacts directly with cyclins and promotes G1 cell cycle arrest by downregulating transcription of cyclin and CDK genes ^41^. Here, we have shown how Pmk1 and Hox7 are both essential for expression of autophagy-related and cell cycle control genes during appressorium morphogenesis. It therefore appears likely that Hox7 is essential to trigger a G1 arrest in the appressorial nucleus following the initial round of mitosis in the germ tube after conidial germination ^7,19^. This may be necessary to repress hyphal-like growth and to trigger conidial cell death, but the mechanism by which this occurs and the precise interplay with the initiation of autophagy in the conidium requires further study. One possibility is that the role of Hox7 is influenced by starvation stress, because *M. oryzae* appressoria only form in free water and their development can be repressed by the presence of exogenous nutrients ^1,36,42^. The nutrient-sensing Snf1 kinase in yeast ^43^, for example, regulates autophagy genes, such as Atg1 and Atg13, acting antagonistically to the cyclin protein kinase Pho85 which in turn, acts as a negative regulator of autophagy ^44^. Moreover, Pho85 has been shown to regulate infection-associated morphogenesis and virulence in pathogenic fungi, such as the corn smut fungus *Ustilago maydis* and the human pathogen *Candida albicans* ^45–47^. In *M. oryzae*, Snf1 mutants are impaired in appressorium development, lipid mobilization and turgor generation ^48^, but the role of Snf1 in control of autophagy and cell cycle progression has not yet been defined. Investigating the interplay of Snf1 and Pho85 with Hox7 may therefore prove valuable in further characterising the Pmk1 signalling pathway.

The role of Mst12, which has long been known to be required for plant infection ^49^, has also been more clearly defined by this study. We were able to show, for instance, that Mst12 regulates appressorium maturation by controlling expression of more than 2000 genes and an additional 53 transcription factors, highlighting the complexity of appressorium maturation and processes involved in cuticle rupture and penetration peg development. Importantly, Mst12 not only regulates expression of genes associated with the re-polarisation process, but also genes associated with plant tissue colonization. A subset of effectors, for example, implicated in suppression of host immunity, require Mst12 for their expression during appressorium maturation at the point of plant infection, as well as genes involved in polarised exocytosis. Taken together, this suggests that early-acting effectors are expressed prior to plant infection in the appressorium and that both their expression and secretion require Mst12.

In summary, we have provided evidence that Pmk1 MAPK acts as a global regulator of appressorium development and fungal invasive growth by controlling a hierarchical network of transcriptional regulators, including Hox7 and Mst12. Future studies will focus on how this network of regulators interact and how they exert spatial and temporal control of gene expression during appressorium-mediated plant infection.

## Materials and Methods

### Fungal Strains, Growth Conditions and plant infection assays

All isolates of *Magnaporthe oryzae* used and generated in this study are stored in the laboratory of N. J. Talbot (The Sainsbury Laboratory). Fungal strains were routinely incubated at 26°C with a 12 h photoperiod. Fungal strains were grown on complete medium (CM) ^1^. Rice infections were performed using a blast-susceptible rice (*Oryza sativa*) cultivar, CO-39 ^50^. For plant infections, conidial suspensions (5×10^4^ conidia ml^-1^ in 0.1% gelatin) were spray inoculation onto 3 week-old seedlings and incubated for 5 days in a controlled environment chamber at 24 °C with a 12 h photoperiod and 90 % relative humidity. Disease lesion density was recorded 5 days post-inoculation, as described previously (Talbot et al., 1993).

### Generation of GFP fusion plasmids and strains expressing GFP and RFP fusions

Corresponding DNA sequences were retrieved from the *M. oryzae* database DNA sequences were retrieved from the *M. oryzae* database (http://fungi.ensembl.org/Magnaporthe_oryzae/Info/Index). In-Fusion cloning (In-Fusion Cloning kit; Clontech Laboratories) was used to generate Flp1–GFP and Flp2–GFP. The primers used are shown in Supplementary Table 1. Amplified fragments were cloned into HindIII-digested 1284 pNEB-Nat-Yeast cloning vector with the *BAR* gene that confers bialophos (BASTA) resistance. Plasmids expressing GFP and RFP fusions were transformed into Guy11, Δ*pmkl*, Δ*mst12* and Δ*hox7* mutants and *M. oryzae* transformants with single-insertions selected by Southern blot analysis. Independent *M. oryzae* transformants were used for screening and selected for consistency of the fluorescence localization.

### Generation of *M.oryzae* targeted gene deletion mutants

Targeted gene replacement mutants of *M. oryzae* (Talbot et al., 1993) were generated using the split marker technique, described previously (Kershaw and Talbot, 2009). Gene-specific split marker constructs were amplified using primers in Supplementary Table 1 and fused with bialophos (BASTA) resistance cassette or hygromycin resistant cassette transformed either into Guy11 or Δ*ku70*. Transformants were selected on glufosinate (30 μg ml^-1^) or hygromycin (200 μg ml^-1^) and assessed by Southern Blot analysis to verify complete deletion of each gene.

### *In vitro* appressorium development assays and live cell imaging

Appressorium development was induced on borosilicate 18×18 glass coverslips, which were termed the HP surface in all experiments (Fisher Scientific UK Ltd). Conidial suspensions were prepared at 5×10^4^ conidia ml^-1^ in double distilled water and 50 μl of the conidial suspension placed onto the coverslip surface and incubated in a controlled environment chamber at 24°C. For incubation on non-inductive surfaces, the hydrophilic surface of Gelbond (Sigma SA) was used and termed the HL surface. Epifluorescence and differential interference contrast (DIC) microscopy was carried out using the IX81 motorized inverted microscope (Olympus, Hamburg, Germany) and images were captured using a Photometrics CoolSNAP HQ2 camera (Roper Scientific, Germany). The system was under the control of MetaMorph software package (MDS Analytical Technologies, Winnersh, UK). Datasets were compared using unpaired Student’s *t*-test.

### RNA extraction and RNA-Seq analysis

Conidia were harvested from 10-day old CM agar plates, washed and conidial suspensions (7.5×10^5^ conidia ml^-1^) prepared in the presence of 50 ng μl^-1^ 1,16-Hexadecanediol (Sigma SA). This spore suspension was poured into square petri plates (Greiner Bio One) to which 10 glass cover slips (Cole-Parmer) had been attached by gluing. Appressorium formation was monitored under a Will-Wetzlar light inverted microscope (Wilovert®, Hund Wetzlar, Germany) for each time point ensuring homogeneous and synchronised infection structure formation. Samples were collected and total RNA extracted using the Qiagen RNeasy Plant Mini kit, according to manufacturer’s instructions. RNA-Seq libraries were prepared using 5 μg of total RNA with True Seq RNA Sample Preparation kit from Illumina (Agilent) according to manufacturer’s instructions, and sequenced using an Illumina 2000 Sequencer. Output short reads were aligned against version 8.0 of *M. oryzae* genome sequence using Tophat software ^51^. Analysis of the data was performed using DESeq which determines differential gene expression through moderated log2 fold change value (mod_lfc) ^52^. Transcript abundances for each gene and adjusted P-values, were generated as described by Soanes *et al.*, (2012). To determine the significant differences of the pair wise comparisons, we adjusted the p-value <=0.01.

### Statistical Analysis

All the experiments were conducted with at least three biological replicates. P-values <0.05 were considered significant; * P-value < 0.05, ** P-value < 0.01, *** P-value < 0.001, **** P-value < 0.0001. P-values >0.05 were considered non-significant and exact values are shown where appropriate. Dot plots were routinely used to show individual data points and generated with Prism7 (GraphPad). Bar graphs showed ± SE and were generated with Prism7 (GraphPad). In pathogenicity assays before comparison, data sets were tested for normal distribution using Shapiro-Wilk normality test. In all cases where at least one data set was non-normally distributed (*P*>0.05 in Shapiro-Wilk tests), we used non-parametric Mann-Whitney testing. Analysis of non-normal data sets are represented by box and shisker blots (=box plots) that show 25/75 percentile the median, and the minimum and maximum values by the ends of the whiskers. For appressorial development assays, data were analysed using unpaired two-tailed Student’s t-testing.

### Protein-protein interaction analysis by yeast two-hybrid assay

Yeast two-hybrid analysis was used to analyse the protein-protein interaction between Pmk1, Mst12 and Hox7 utilizing the Matchmaker Gold System (Takara Bio, USA). Fragments of Pmk1, Mst12 and Hox7 were amplified from cDNA derived from lyophilized *M. oryzae* appressoria generated on borosilicate 18 × 18-mm glass coverslips (Thermo Fisher Scientific) using the primers of Supplementary Table 1. Fragments were cloned in both pGBKT7 (bait) and pGADT7 (prey) vectors by In-fusion cloning (In-Fusion Cloning kit; Clontech Laboratories). BKT variants and GAD were co-transformed into chemically competent Y2HGold yeast cells (Takara Bio, USA) following the manufacturer’s instructions.

For each interaction tested, single colonies were grown on SD-Leu-Trp selection media at 30°C. Successfully transformed yeast colonies were inoculated in 5ml of liquid SD-Leu-Trp selection media for overnight growth at 30°C. The liquid culture was then used to make serial dilutions of OD600 1, 1^-1^, 1^-2^, and 1^-3^, respectively. As a control, 5μl droplets of each dilution were spotted on a SD-Leu-Trp plate and to detect the interactions, they were spotted on a SD-Leu-Trp-Ade-His plate containing X-a-gal and Aureobasidin A antibiotic (Takara Bio, USA). After incubation for 48-72 hours at 30°C, the plates were analysed and imaged. All experiments were performed in triplicate.

### Proteinextraction, phosphoprotein-enrichment, mass spectrometry analysis for Discovery Phosphoproteomics, Parallel Reaction Monitoring and Phospho-peptide quantitation

Total protein was extracted from lyophilized *M. oryzae* appressoria from Guy11 and germlings of *Δpmkl* mutants generated on borosilicate 18 × 18-mm glass coverslips (Thermo Fisher Scientific) at 0, 1, 1.5 and 2 h for the Parallel Reaction Monitoring (PRM) experiment. For discovery phosphoproteomics, total protein was extracted from lyophilized *M. oryzae* appressoria from Guy11, *Δpmkl* and *Δhox7* mutants generated on borosilicate 18 × 18-mm glass coverslips (HP) (Thermo Fisher Scientific) at 6h, as performed for the RNA-seq experiements. Lyophilized appressoria were resuspended in extraction buffer (8 M urea,150 mM NaCl, 100 mM Tris pH 8, 5 Mm EDTA, aprotinin 1 μg/mL, leupeptin 2 μg/mL) and mechanically disrupted in a 2010 GenoGrinder® tissue homogenizer (1 min at 1300 rpm). The homogenate was centrifuged for 10 min at 16,000 × *g*, at 4 °C (Eppendorf Micro-centrifuge 5418). The supernatant was removed and used to determine total protein concentration using the Bradford Assay. For phosphopeptide enrichment, sample preparation started from 1-3 mg of total protein extract dissolved in ammonium bicarbonate buffer containing 8 M urea. First, protein extracts were reduced with 5 mM Tris (2-carboxyethyl) phosphine (TCEP) for 30 min at 30°C with gentle shaking, followed by alkylation of cysteine residues with 40mM iodoacetamide at room temperature for 1 hour. Samples were diluted to a final concentration of 1.6 M urea with 50mM ammonium bicarbonate and digested overnight with trypsin (Promega; 1:100 enzyme to substrate ratio). Peptide digests were purified using C18 SepPak columns (Waters), as described previously ^53^. Phosphopeptides were enriched using titanium dioxide (TiO2, GL Science) with phthalic acid as a modifier ^53^. Finally, phosphopeptides were eluted by a pH-shift to 10.5 and immediately purified using C18 microspin columns (The Nest Group Inc., 5 – 60 μg loading capacity). After purification, all samples were desiccated in a speed-vac, stored at −80°C and re-suspended in 2% Acetonitrile (AcN) with 0.1% trifluoroacetic acid (TFA) before mass-spectrometry analysis ^54^.

LC-MS/MS analysis was performed using an Orbitrap Fusion trihybrid mass spectrometer (Thermo Scientific) and a nanoflow ultra-high-performance liquid chromatography (UHPLC) system (Dionex Ultimate3000, Thermo Scientific). Peptides were trapped to a reverse phase trap column (Acclaim PepMap, C18 5 μm, 100 μm x 2 cm, Thermo Scientific). Peptides were eluted in a gradient of 3-40 % acetonitrile in 0.1 % formic (solvent B) acid over 120 min followed by gradient of 40-80 % B over 6 min at a flow rate of 200 nL/min at 40°C. The mass spectrometer was operated in positive-ion mode with nano-electrospray ion source with ID 0.02mm fused silica emitter (New Objective). A voltage of 2,200 V was applied via platinum wire held in PEEK T-shaped coupling union with transfer capillary temperature set to 275 °C. The Orbitrap mass spectrometry scan resolution of 120,000 at 400 m/z, range 300-1,800 m/z was used, and automatic gain control was set to 2×10^5^ and the maximum inject time to 50 ms. In the linear ion trap, MS/MS spectra were triggered using a data-dependent acquisition method, with ‘top speed’ and ‘most intense ion’ settings. The selected precursor ions were fragmented sequentially in both the ion trap using collision-induced dissociation (CID) and in the higher-energy collisional dissociation (HCD) cell. Dynamic exclusion was set to 15 sec. The charge state allowed between 2+ and 7+ charge states to be selected for MS/MS fragmentation.

Peak lists in format of Mascot generic files (.mgf files) were prepared from raw data using MSConvert package (Matrix Science). Peak lists were searched on Mascot server v.2.4.1 (Matrix Science) against either *Magnaporthe oryzae* (isolate 70-15, version 8) database, an in-house contaminants database. Tryptic peptides with up to 2 possible miscleavages and charge states +2, +3, +4, were allowed in the search. The following modifications were included in the search: oxidized methionine, phosphorylation on Serine, Threonine, Tyrosine as variable modification and carbamidomethylated cysteine as static modification. Data were searched with a monoisotopic precursor and fragment ions mass tolerance 10ppm and 0.6 Da respectively. Mascot results were combined in Scaffold v. 4 (Proteome Software) to validate MS/MS-based peptide and protein identifications and annotate spectra. The position of the modified residue and the quality of spectra for individual phosphopeptides were manually inspected and validated.

Peptide quantitation was performed using Parallel Reaction Monitoring (PRM) as described previously^55^. Briefly, phospho-peptide DNRPLS[+80]PVTLSGGPR was targeted to measure Hox7 phosphorylation at serine 158. The PRM assay also included a selection of control peptides (Supplementary Table x) having similar relative intensities in each sample and used to measure relative phospho-peptide content. DNRPLS[+80]PVTLSGGPR intensity was normalised against the summed control peptide intensities to correct for differences in phospho-peptide yield. The assay was performed once for each of three biological replicates and results averaged ± SE.

## Acknowledgements

This project was supported by a European Research Council Advanced Investigator award (to N.J.T.) under the European Union’s Seventh Framework Programme FP7/2007-2013/ERC Grant Agreement 294702 GENBLAST and by the Gatsby Charitable Foundation.

## Author Contributions

M.O-R. and NJT conceptualized the project. Experimental analyses were carried out by M.O-R, M.M-U., M.J.K., and C.M. N.C-M. P.D. and F.L.H.M carried out phosphoproteomic analysis. G.V.P and B.V. generated the *Drpp3* mutant. Bioinformatic analysis was performed by M.O-R. and D.M.S. The paper was written by M.O-R. and NJT, with contributions from all authors.

## Competing Interests

The authors declare no competing interests

## Data availability

All data that support the findings of this paper are available from the corresponding author on request. RNA-seq data described in this paper has been submitted to European Nucleotide Archive ENA https://www.ebi.ac.uk/ena/submit, under accession number PRJEB36580 and study unique number ena-STUDY-JIC-04-02-2020-15:54:49:677-189. All *Magnaporthe oryzae* strains generated in this study are freely available upon request from the corresponding author.

