## Supplementary Figures for "A hierarchical transcriptional network controls appressorium-mediated plant infection by the rice blast fungus *Magnaporthe oryzae*"

**a**

| Name | Accession | SignalP_5.0 | |
|---|---|---|---|
| | | Probability | Position |
| ***FLP1*** | MGG_02884 | 05391 | 17-18 |
| ***FLP2*** | MGG_09372 | 0.8169 | 16-17 |
| ***FLP3*** | MGG_05483 | 0.5644 | 18-19 |

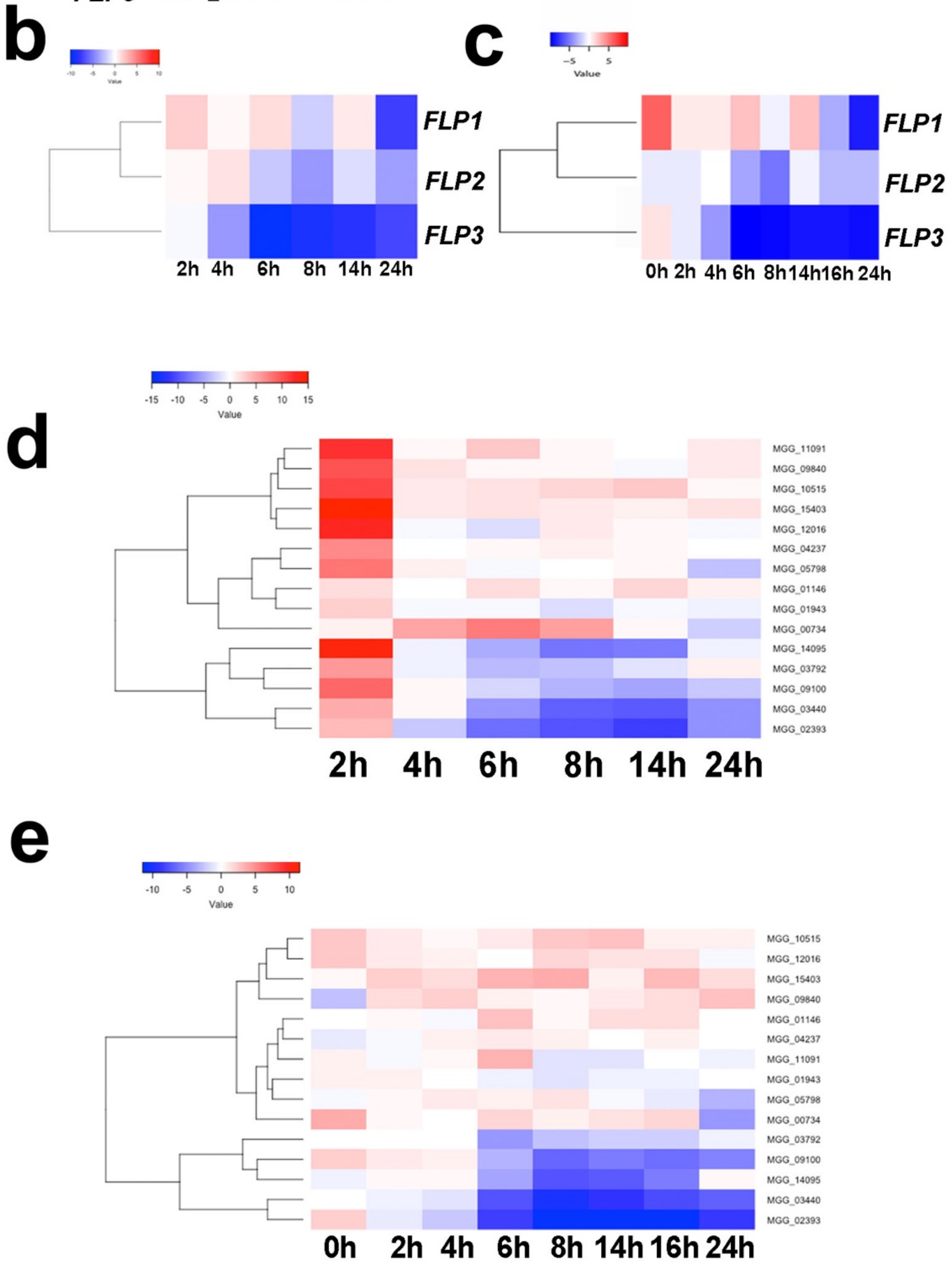


**Supplementary figure 1: Gene expression of fasciclin-related (PF02469) and cutinase-encoding genes (PF01083) in *M. oryzae* occurs in response to HP surfaces and is Pmk1-dependent.** a. Table to show the probability and position of signal peptide prediction for Flp1, Flp2 and Flp3 in *M. oryzae* using http://www.cbs.dtu.dk/services/SignalP-5.0/. b. Heatmap to show relative patterns of transcript abundance of fasciclin-encoding genes in Guy11 in response to HL *vs.* HP surfaces between 0h and 24h after conidial germination. c. Heatmap to show relative patterns of transcript abundance of fasciclin encoding genes in *Δpmk1* mutant *vs.* Guy11 following incubation on HP surfaces between 0h and 24h after conidial germination. d. Heatmap to show relative levels of transcript abundance of 15 putative cutinase-encoding genes in Guy11 in response to HL *vs.* HP surfaces between 0h and 24h after conidial germination e. Heatmap to show relative levels of 15 putative cutinase-encoding genes in *Δpmk1* mutant *vs.* Guy11 following incubation on HP surface between 0h and 24h after conidial germination.

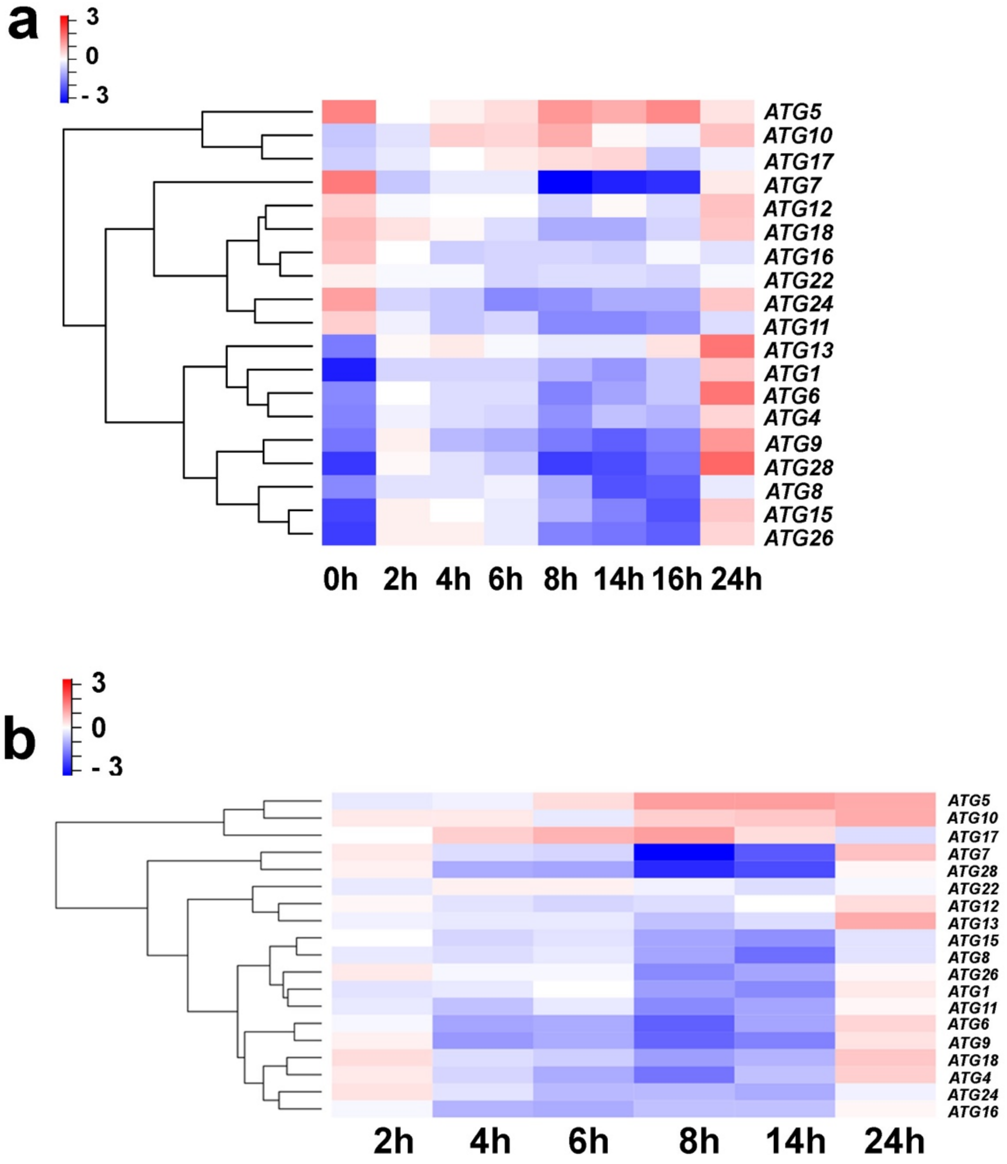


**Supplementary figure 2: Gene expression of autophagy-related genes in *M. oryzae* occurs in response to HP surfaces and is Pmk1-dependent.** a. Heatmap to show relative levels of transcript abundance of 19 autophagy-related genes in a *Δpmk1* mutant *vs.* Guy11 following incubation on HP surfaces between 0h and 24h after conidial germination. b. Heatmap to show relative levels of transcript abundance of 19 autophagy-related genes in Guy11 in response to HL *vs.* HP surfaces between 0h and 24h after conidial germination.

**a**

| Cell cycle-related genes | | |
|---|---|---|
| **Information related** | **Gene Name** | **Accesion Number** |
| Putative STE5 homologue | *FAR1* | MGG_00134 |
| DNA damage checkpoint regulator | *DBF4/NIM1* | MGG_00597 |
| CDK regulatory subunit | *CKS1* | MGG_00682 |
| Mitotic Phosphatase | *CDC14* | MGG_00757 |
| Mitotic Phosphatase | *TINA* | MGG_00763 |
| DDR pathway-related protein | *DUN1* | MGG_01596 |
| Mitotic CDK | *CDC28* | MGG_01362 |
| CDK kinase | *SWE1* | MGG_01816 |
| Nim1-related kinase | *GIN4* | MGG_02810 |
| Serine/ Threonine protein kinase | *NIMA* | MGG_03026 |
| APC/C subunit | *APC1/BIM1* | MGG_03314 |
| G1-type cyclin | *CLN3* | MGG_03595 |
| DDR pathway-related protein | *CHK1* | MGG_03729 |
| APC/C subunit | *APC2* | MGG_04724 |
| DDR pathway-related protein | *CDS1* | MGG_04790 |
| B-type cyclin | *CYC1* | MGG_05646 |
| B-type cyclin | *CYC2* | MGG_07065 |
| CDK phosphatase | *MIH1* | MGG_07734 |
| Polo kinase | *CDC5* | MGG_09960 |
| CDK7-like CDK | *KIN28* | MGG_13401 |
| Nim1-related kinase | *HSL7* | MGG_03894 |

**b**

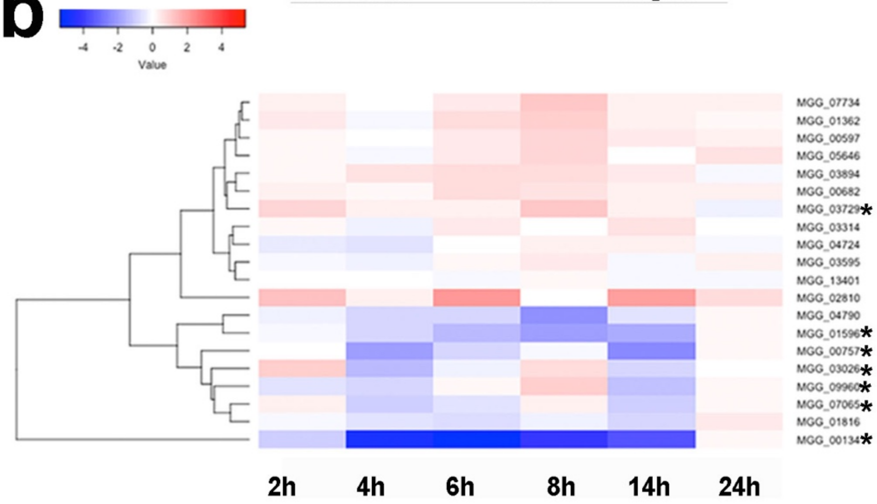


**c**

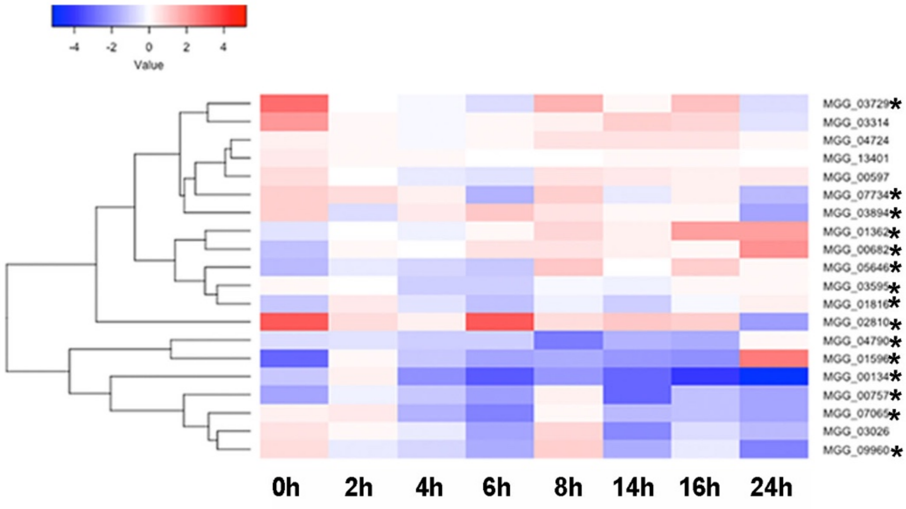


**Supplementary figure 3: Cell cycle related genes are expressed in a Pmk1 dependent manner during appressorium development by *M. oryzae*.** a. Table to show 21 putative cell cycle-related genes selected for transcriptomic analysis [1]. b. Heatmap to show relative levels of transcript abundance of cell cycle-associated genes in Guy11 in response to HL *vs.* HP surfaces between 0h and 24h after conidial germination. c. Heatmap to show relative levels of transcript abundance of cell cycle associated genes in Δ*pmk1* mutant vs. Guy11 following incubation on HP surfaces between 0h and 24h after conidial germination. Asterisks (*) indicate differentially-regulated genes (padj< 0.01) in at least one time point during appressorium development by *M. oryzae*.

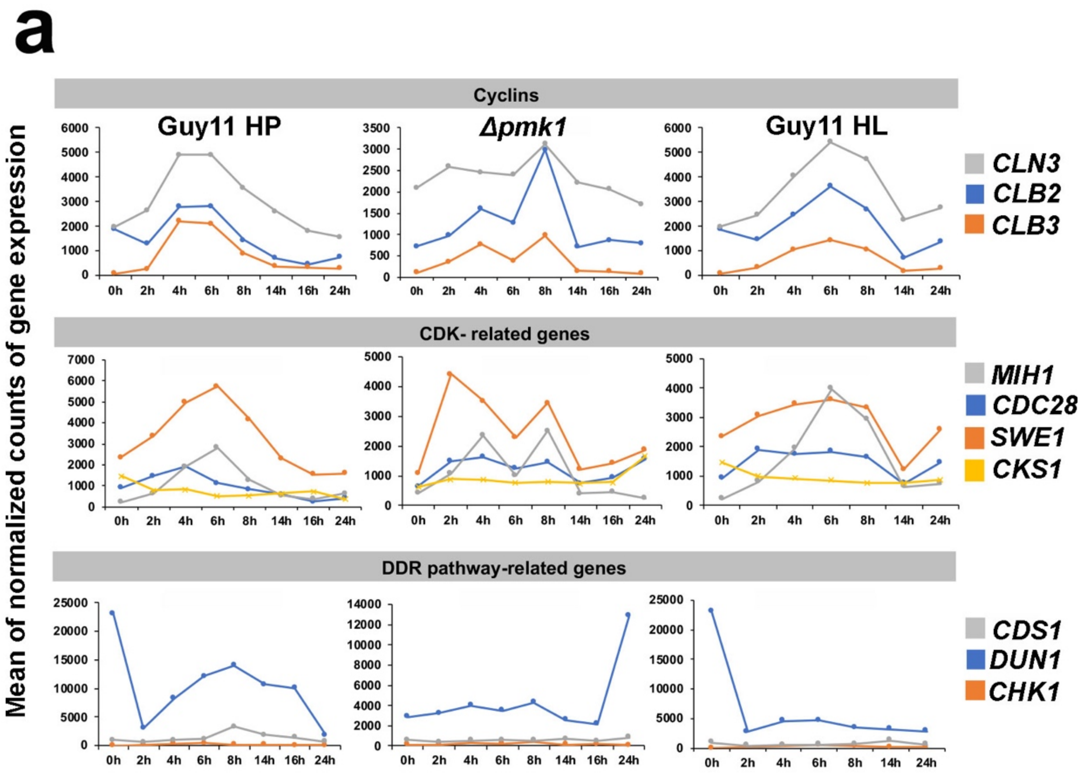


**Supplementary figure 4: Temporal expression profiles of cell cycle related genes in *M. oryzae* in response to HP surfaces and the presence of the Pmk1 MAPK.** a. Line graphs to show mean of normalized counts of gene expression of cyclins (*CLN3, CLB2* and *CLB3*), Cdk-related genes (*MIH1, CDC28, SWE1* and *CKS1*) and DNA Damage Response (DDR) pathway-related genes (*CDS1, DUN1* and *CHK1*) during a time course of appressorium development in Guy11 on HP surface, the *Δpmk1* mutant on an HP surface and Guy11 on HL surface between 0h and 24h after conidial germination.

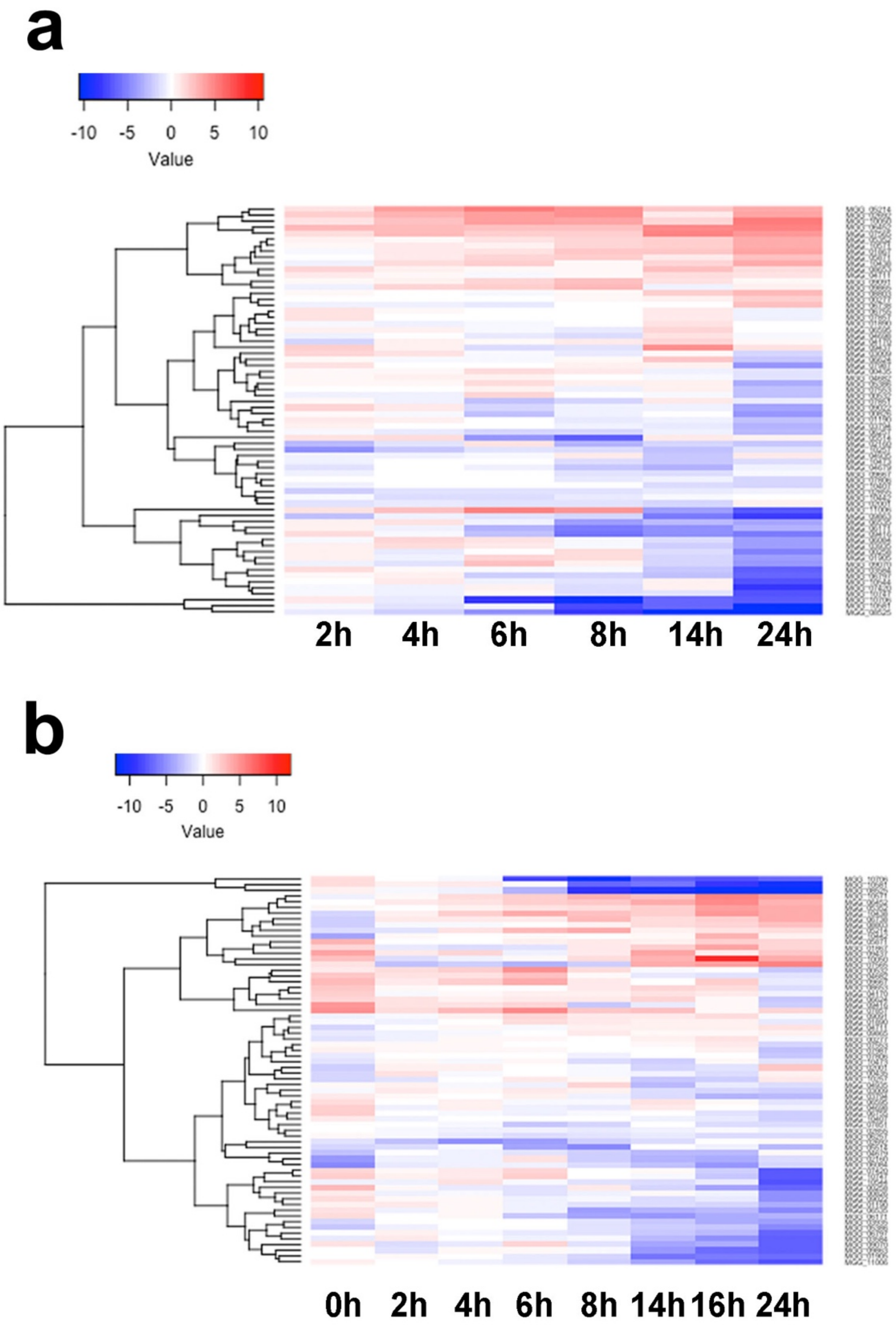


**Supplementary figure 5: Gene expression of putative G-protein coupled receptor-encoding genes in *M. oryzae* in response to HP surface and the presence of the Pmk1 MAP kinase.** a. Heatmap to show relative in response to HL *vs.* HP surfaces between 0h and 24h after conidial germination. b. Heatmap to show relative levels of transcript abundance of 79 putative G-coupled receptor encoding genes in *M. oryzae* in a Δ*pmk1* mutant vs. Guy11 on HP surfaces between 0h and 24h following conidial germination.

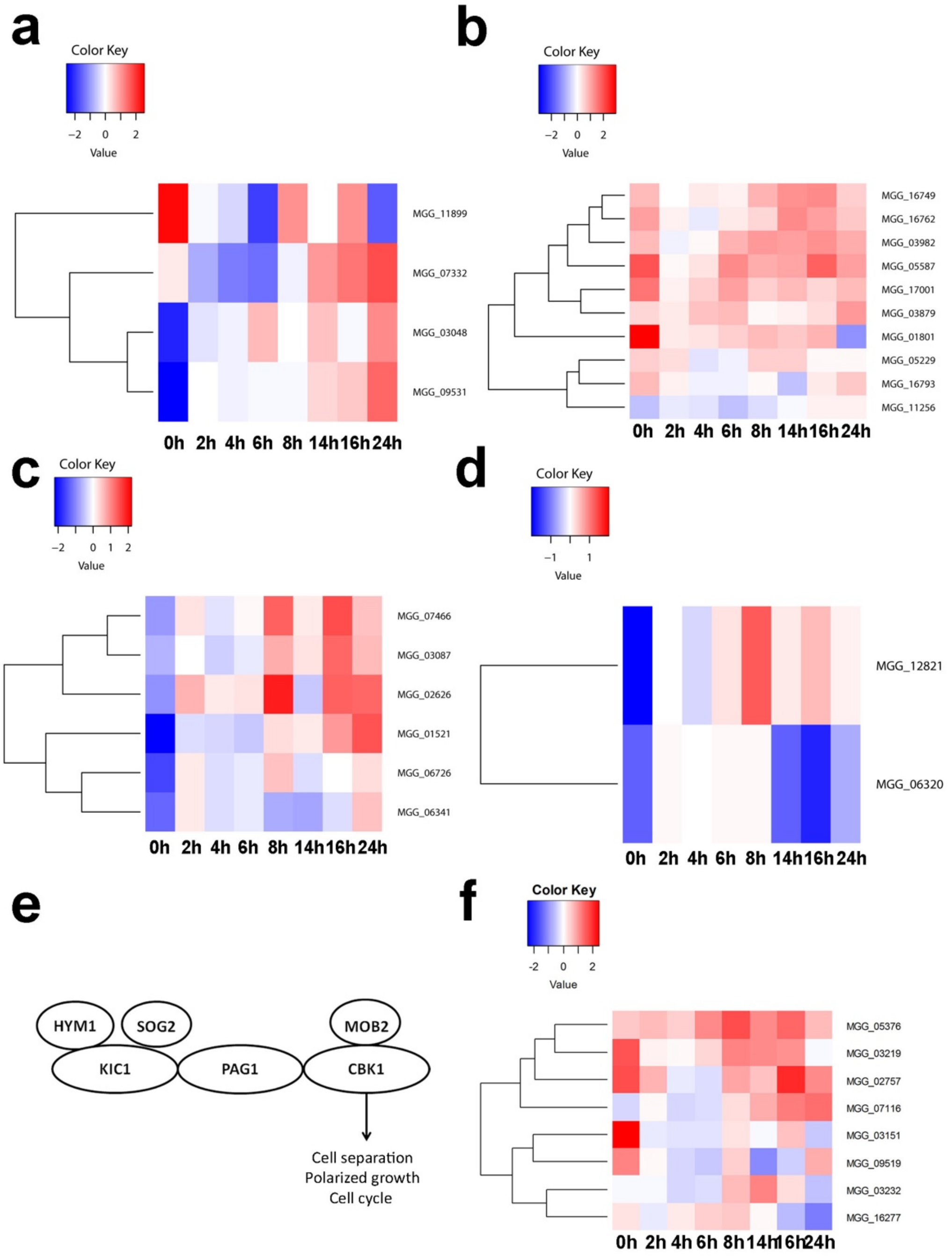


**Supplementary figure 6: Expression of cytoskeleton-re-modelling, septin GTPase and RAM pathway-related genes is dependent on the Pmk1 MAP kinase during appressorium development by *M. oryzae*.** a. Heatmap to show relative levels of transcript abundance of Actin (PF00022)-domain containing genes in a Δ*pmk1* mutant vs.

Guy11 on HP surface between 0h and 24h following conidial germination. b. Heatmap to show relative levels of transcript abundance of Fes/CIP4, and EFC/F-BAR homology domain (PF00611)-domain containing genes in a Δ*pmk1* mutant vs. Guy11 on HP surface between 0h and 24h following conidial germination. c. Heatmap to show relative levels of transcript abundance of Septin (PF00735)-domain containing genes in a Δ*pmk1* mutant vs. Guy11 on HP surface between 0h and 24h following conidial germination. d. Heatmap to show relative levels of transcript abundance of P21-Rho-binding domain (PF00786) and protein kinase domain (PF00069)-domain containing genes a Δ*pmk1* mutant vs. Guy11 on HP surface between 0h and 24h following conidial germination. e. Model to show the components of a putative *M. oryzae* RAM pathway according, to a model of the human pathogen *C. albicans.* f. Heatmap to show relative levels of transcript abundance of RAM pathway-associated genes differentially regulated in a Δ*pmk1* mutant vs. Guy11 on HP surface between 0h and 24h following conidial germination. (blue= down-regulated in Δ*pmk1* mutant vs. Guy11; red= up-regulated in Δ*pmk1* mutant vs. Guy11).

**a**

| Gene Name | # Accession |
|---|---|
| Acetyl-CoA carboxylase | MGG_07613 |
| Polyketide synthase (*ALB1*) | MGG_07219 |
| Trahydroxynaphthalene reductase (*T4HN*) | MGG_07216 |
| Scytalone degydratase (*RSY1*) | MGG_05059 |
| Trihydroxynaphthalene reductase (*BUF1*) | MGG_02252 |

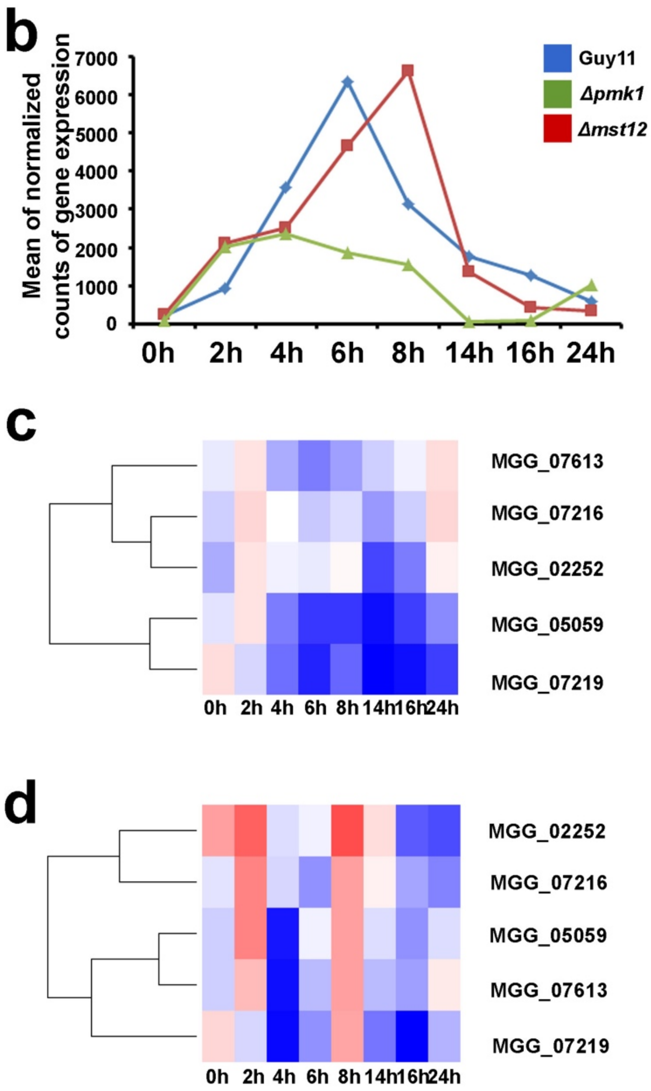


**Supplementary figure 7: Expression of melanin biosynthesis pathway genes during appressorium development by *M. oryzae*.** a. Table to show names and accession numbers of melanin biosynthesis enzymes in *M. oryzae*. b. Graph to show mean of normalized counts of gene expression of melanin biosynthesis enzymes during appressorium development time course for Guy11 wild type strain, *Δpmk1* mutant and *Δmst12* mutant on HP surface between 0h and 24h following conidial germination. c. Heatmap showing levels of transcript abundance from genes involved in melanin biosynthesis in a Δ*pmk1* mutant compared to Guy11 HP surface between 0h and 24h following conidial germination. d. Heatmap showing levels of transcript abundance of genes involved in melanin biosynthesis in a Δ*mst12* mutant compared to Guy11 HP surface between 0h and 24h following conidial germination. Levels of expression are represented as moderated logarithmic fold change (mod_lfc.  (blue= down-regulated in mutants vs. Guy11; red= up-regulated in mutants vs. Guy11).

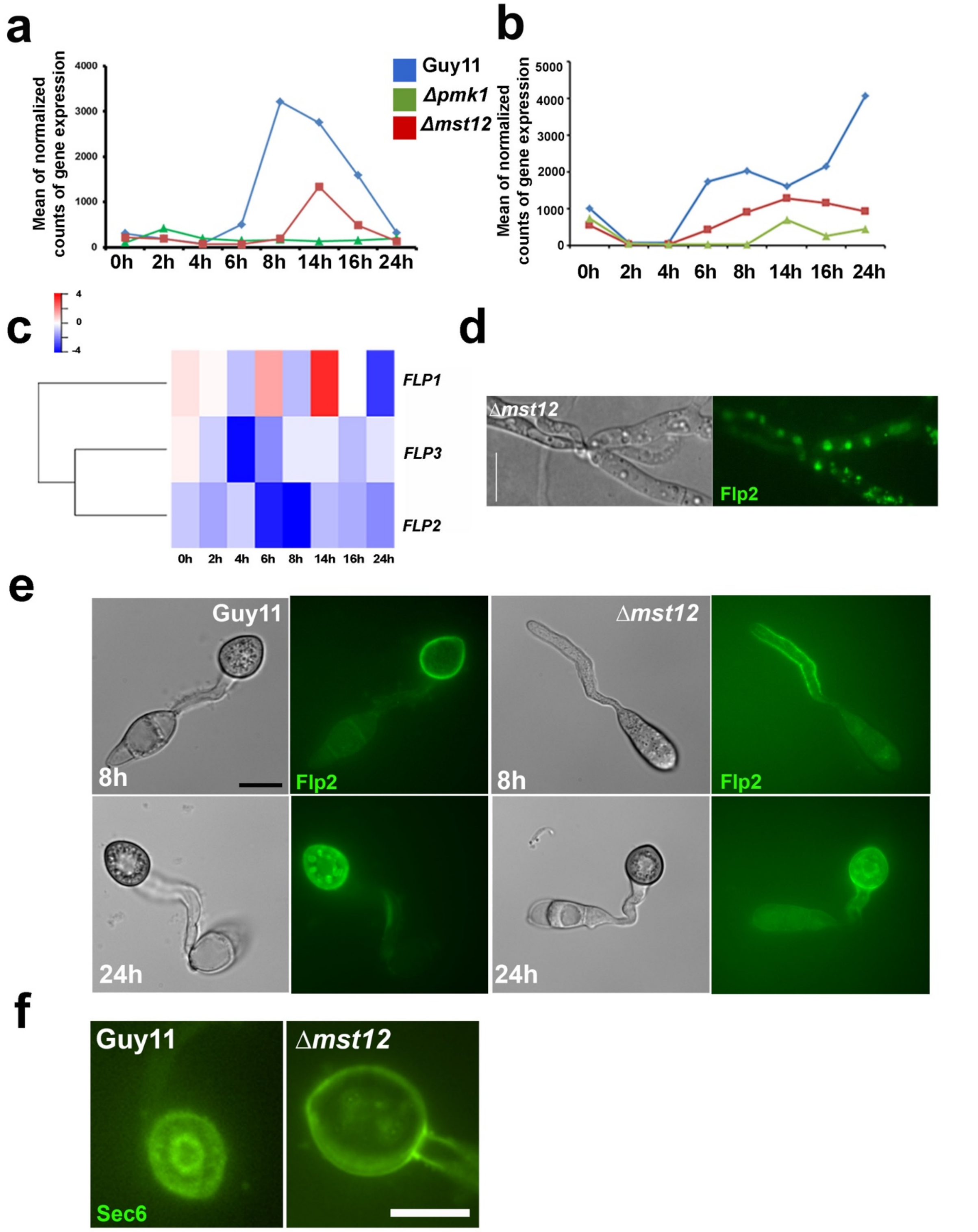


**Supplementary figure 8: Cutinase and fasciclin family gene expression and sub-cellular localization is dependent on Mst12 during appressorium development by *M. oryzae*.** a. Line graph to show mean normalized counts of gene expression of cutinase-encoding genes during a time course of appressorium development by

Guy11, *Δpmk1* and *Δmst12* mutants on HP surface between 0h and 24h. b. Line graph to show mean normalized counts of gene expression of fasciclin-encoding genes in Guy11, *Δpmk1* and *Δmst12* mutants on HP surface between 0h and 24h. c. Heatmap showing levels of transcript abundance of fasciclin genes in a *Δmst12* mutant compared to Guy11 during appressorium formation. (blue= down-regulated in mutant vs. Guy11; red= up-regulated in mutant *vs.* Guy11). d. Light micrograph to show sub-cellular localization of Flp2-GFP during live cell imaging of mycelium of a *Δmst12* mutant. Bright field image in left hand panel. Epifluorescence image on right. Bar = 10µm e. Light micrographs to show Flp2-GFP localization during appressorium development in Guy11 and a *Δmst12* mutant. Bright field image in left hand panels. Epifluorescence images on right. Bar for all panels = 10µm f. Light micrographs to show localization of Sec6-GFP during appressorium development in Guy11 and *Δmst12* mutant. Bright field image in left hand panel. Epifluorescence image on right. Bar = 10µm.

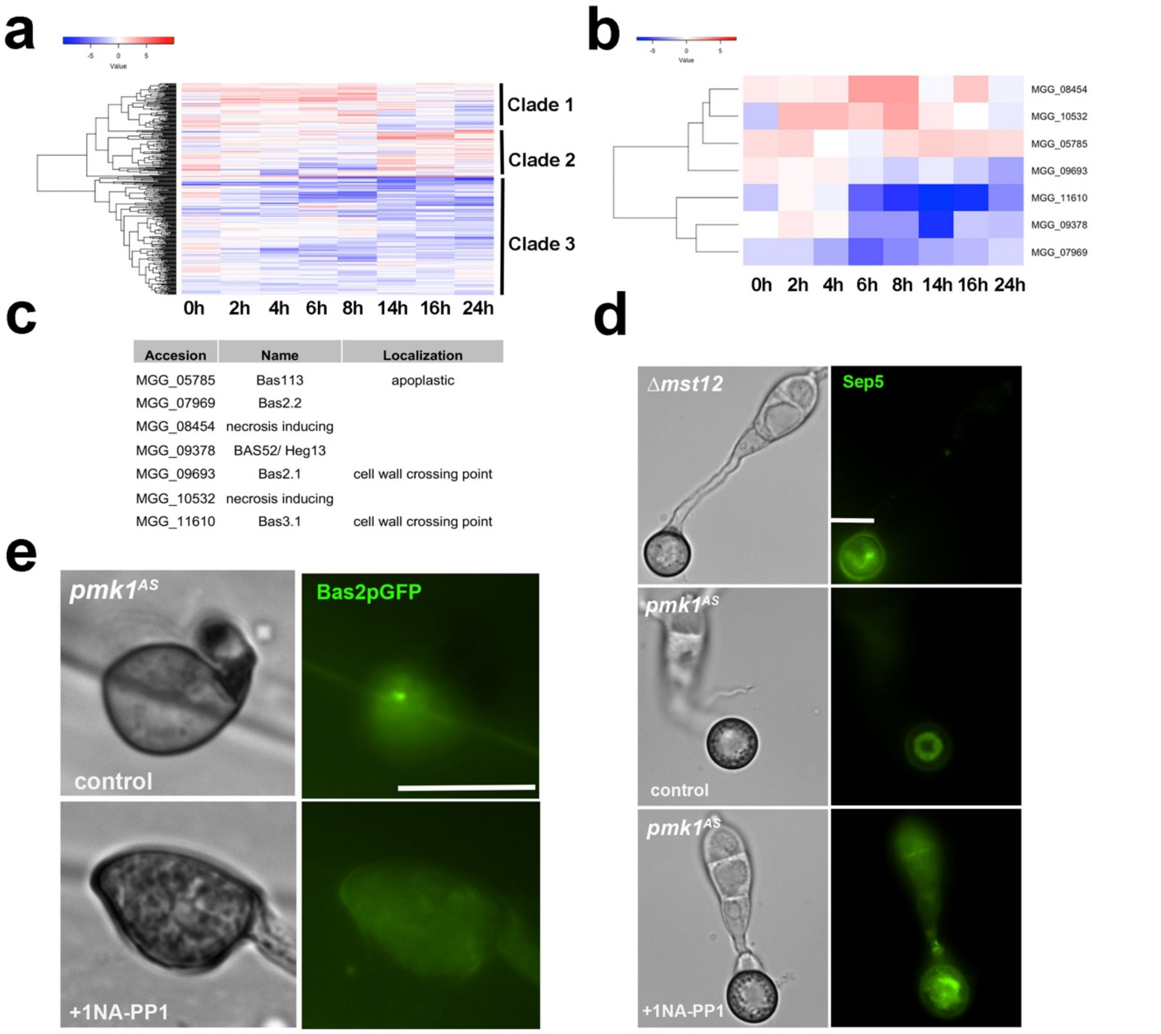


| Accesion | Name | Localization |
|---|---|---|
| MGG_05785 | Bas113 | apoplastic |
| MGG_07969 | Bas2.2 | |
| MGG_08454 | necrosis inducing | |
| MGG_09378 | BAS52/ Heg13 | |
| MGG_09693 | Bas2.1 | cell wall crossing point |
| MGG_10532 | necrosis inducing | |
| MGG_11610 | Bas3.1 | cell wall crossing point |

**Supplementary figure 9: Mst12-dependent gene expression and sub-cellular localisation of a sub-set of *M. oryzae* effectors during appressorium development.** a. Heatmap showing temporal pattern of transcript abundance of 436 genes encoding secreted proteins during appressorium development between 0h and 24h on a HP surface. All were Mst12-dependent and associated with appressorium maturation. b. Heatmap showing levels of transcript abundance of 7 known effector-encoding genes [2] in a Δ*mst12* mutant during appressorium development. (blue= down-regulated in Δ*mst12* mutant vs. Guy11; red= up-regulated in Δ*mst12* mutant vs. Guy11). c. Table to show name and accession number of 7 known effectors differentially regulated in a Δ*mst12* mutant compared to Guy11*.* d. Live cell imaging of Septin Sep5-GFP expression in a Δ*mst12* mutant and $pmk1^{as}$ mutant [3] on HP surface, 24h following conidial germination. The Pmk1asmutant was incubated in the presence or absence of 5µM Na-PP1 added at 6-8h. The Na-PP1 inhibitor is a specific inhibitor of the analogue-sensitive version of Pmk1 expressed by this mutant [3]. e. Micrographs to show expression of the effector Bas2p-GFP in a $pmk1^{as}$ mutant during appressorium formation, 24h after conidial germination inoculated on rice leaf sheath of blast-susceptible rice cultivar Mokoto, in the presence or absence of 5µM Na-PP1 inhibitor. Bar = 10µm.

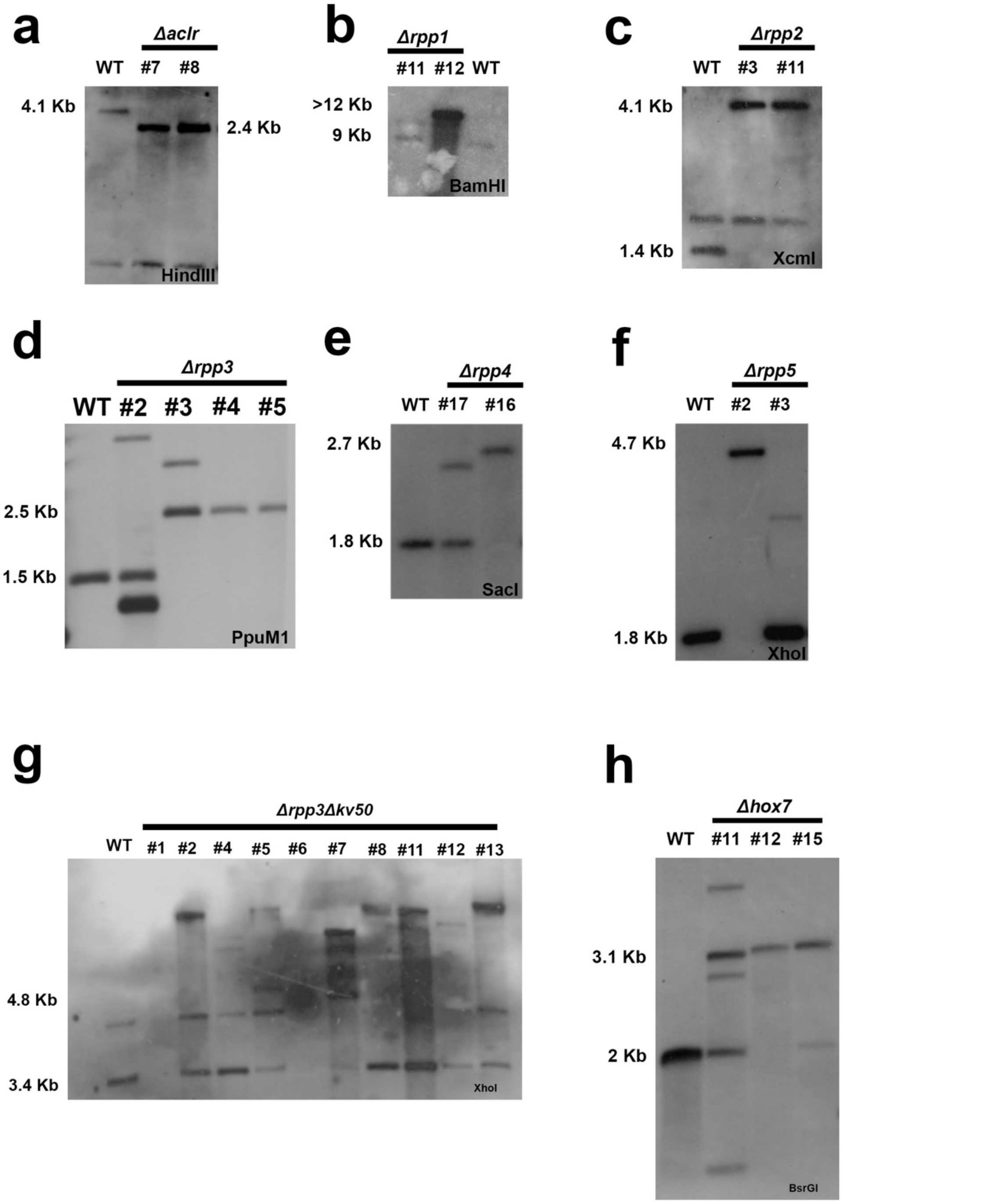


**Supplementary figure 10: Southern Blot analysis to verify generation of targeted gene replacement mutants of clade 4-associated TF-encoding genes in *M. oryzae*** a. Southern blot analysis of genomic DNA extracted from two *M. oryzae* transformants #7 and #8, digested with *Hin*dIII and fractionated by gel electrophoresis. The Southern blot was probed with a 1 kb upstream region of *ACLR* which identified a 4.1 kb restriction fragment when no gene replacement had occurred and a 2.4 kb fragment when the corresponding bialaphos resistance gene cassette was integrated at the *ACLR* locus. b. Southern blot analysis analysis of genomic DNA extracted from two *M. oryzae* transformants #11 and #12 digested with *BamH*I, fractionated by gel electrophoresis and probed with 1 kb upstream region of *RPP1* which identified a 9 kb restriction when no gene replacement had occurred and a >12 kb fragment when the corresponding resistance cassette was integrated at the *RPP1* locus. Southern blot analysis showed a positive result for #12. c. Southern blot analysis of genomic DNA extracted from two *M. oryzae* transformants #3 and #11, digested with *Xmn*I, fractionated by gel electrophoresis and probed with 1 kb upstream region of *RPP2* which identified a 1.4 kb restriction fragment when no gene replacement had occurred and a 4.1 kb fragment when the corresponding resistance cassette was integrated at the *RPP2* locus. Southern blot analysis showed a positive result for both #3 and #11. d. Southern blot analysis of genomic DNA extracted from *M. oryzae* transformants, #2, #3, #4 and #5, digested with *PpuMI*, fractionated by gel electrophoresis and probed with a 796 bp downstream region of *RPP3* which revealed a 1.5 kb restriction fragment when no gene replacement had occurred and a 2.5 kb fragment when the corresponding resistance cassette was integrated at the *RPP3* locus. Southern blot analysis showed a positive result for transformants #4 and #5. e. Southern blot analysis of genomic DNA extracted from *M. oryzae* transformants #7 and #10 digested with *Sac*I, fractionated by gel electrophoresis and probed with a 1 kb upstream region of *RPP4* which identified a 1.8 kb restriction fragment when no gene replacement had occurred and a 2.7 kb band when the corresponding resistance cassette was integrated at the *RPP4* locus. Southern blot analysis showed a positive result for #16. f. Southern blot analysis of genomic DNA extracted from *M. oryzae* transformants #2 and #3, digested with *Xho*I, fractionated by gel electrophoresis and probed with a 1 kb upstream region of *RPP5* which identified a 1.8 kb restriction fragment when no gene replacement had occurred and a 4.7 kb band when the corresponding resistance cassette was integrated at the *RPP5* locus. Southern blot analysis showed a positive result for #2. g) Southern blot analysis of genomic DNA extracted from *M. oryzae* transformants #1, #2, #4, #5, #6, #7, #8, #11, #12, #13 of *Δrpp3* mutant genomic DNA extracted from *M. oryzae* transformants mutant, digested with *Xho*I, fractionated by gel electrophoresis, and probed with a 1 kb upstream region of *PIG1* which identified a 3.4 kb restriction fragment when no gene replacement had occurred and a 4.8 kb fragment when the corresponding resistance cassette was integrated at the *PIG1* locus. Southern blot analysis showed a #7 was a *Δrpp3 Δpig1* double mutant h) Southern blot analysis of genomic DNA extracted from *M. oryzae* transformants #11, #12, and #15, digested with *Bsr*GI and probed with a 1 kb upstream region of *HOX7* that identified a 2 kb restriction fragment when no gene replacement had occurred and a 3.1 kb fragment when the corresponding resistance cassette was integrated at the *HOX7* locus. Southern blot analysis showed #12 was a *Δhox7* mutant.

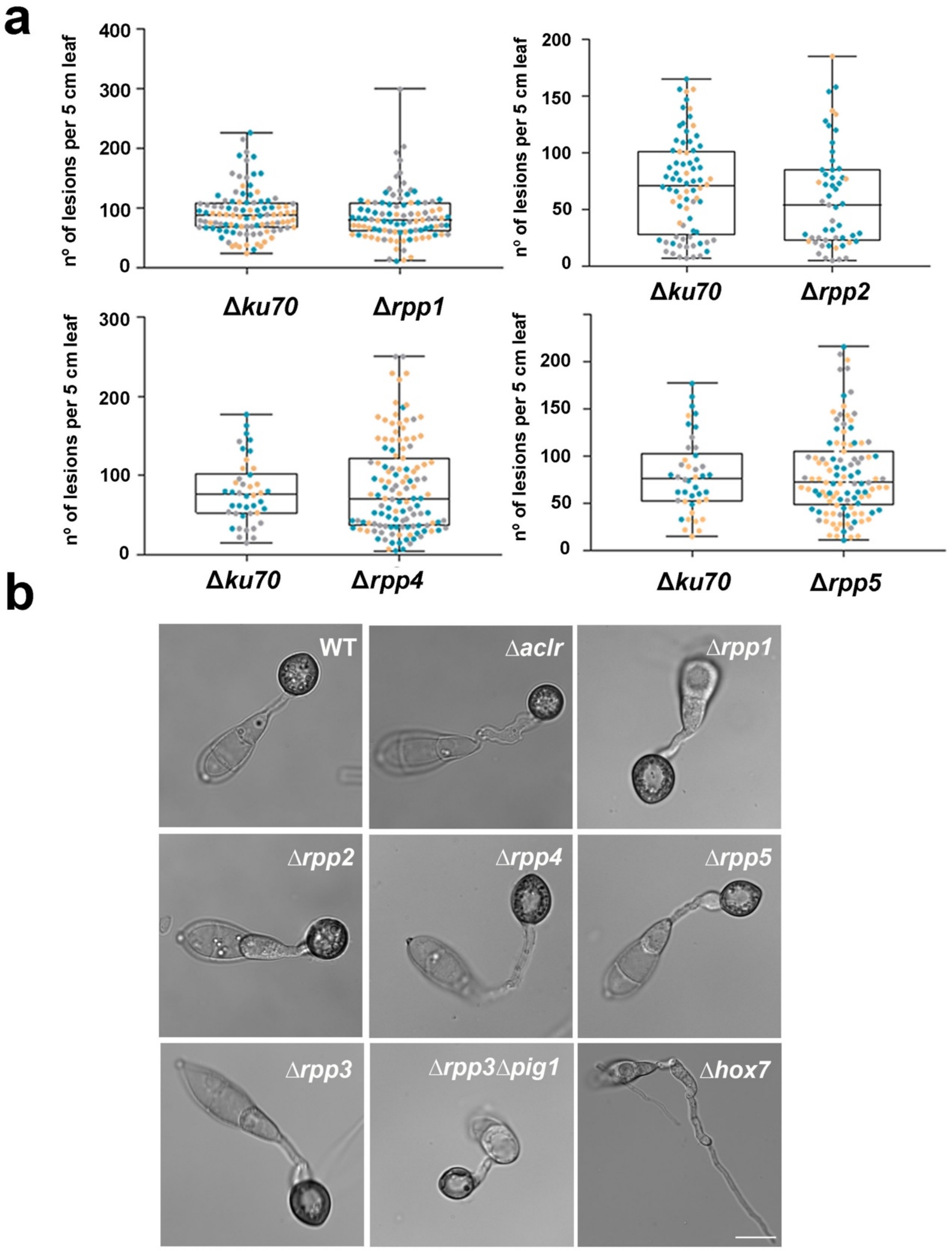


**Supplementary figure 11: Appressorium development and pathogenicity assay of clade 4-associated transcription factor null mutants.** a. Box and Whisker plots to show rice blast disease lesion density per 5m of leaf tissue. 21-day-old rice seedlings of cultivar Co-39 were inoculated with uniform conidial suspensions (5 x $10^4$ conidia $ml^{-1}$) of *Δrpp1, Δrpp2, Δrpp4* and *Δrpp5* mutant and the isogenic wild type Guy11. The plots show the results from 3 biological replications of the experiment and data points are colour-coded for each replication. Error bar shows the standard deviation. b. Light micrographs showing appressorium development of Guy11,

*ΔaclR, Δrpp1, Δrpp2, Δrpp4, Δrpp5, Δrpp3, Δrpp3Δpig1* and *Δhox7* mutants germinated on HP surfaces and observed 24h following conidial germination. Scale bar = 10µm for all panels.

**a**

| 14h | Up mlfc>1 | Down mlfc<-1 | Total |
|---|---|---|---|
| *Δpmk1* | 1636 | 2591 | 4227 |
| *Δmst12* | 632 | 866 | 1498 |
| *Δhox7* | 1879 | 2332 | 4211 |

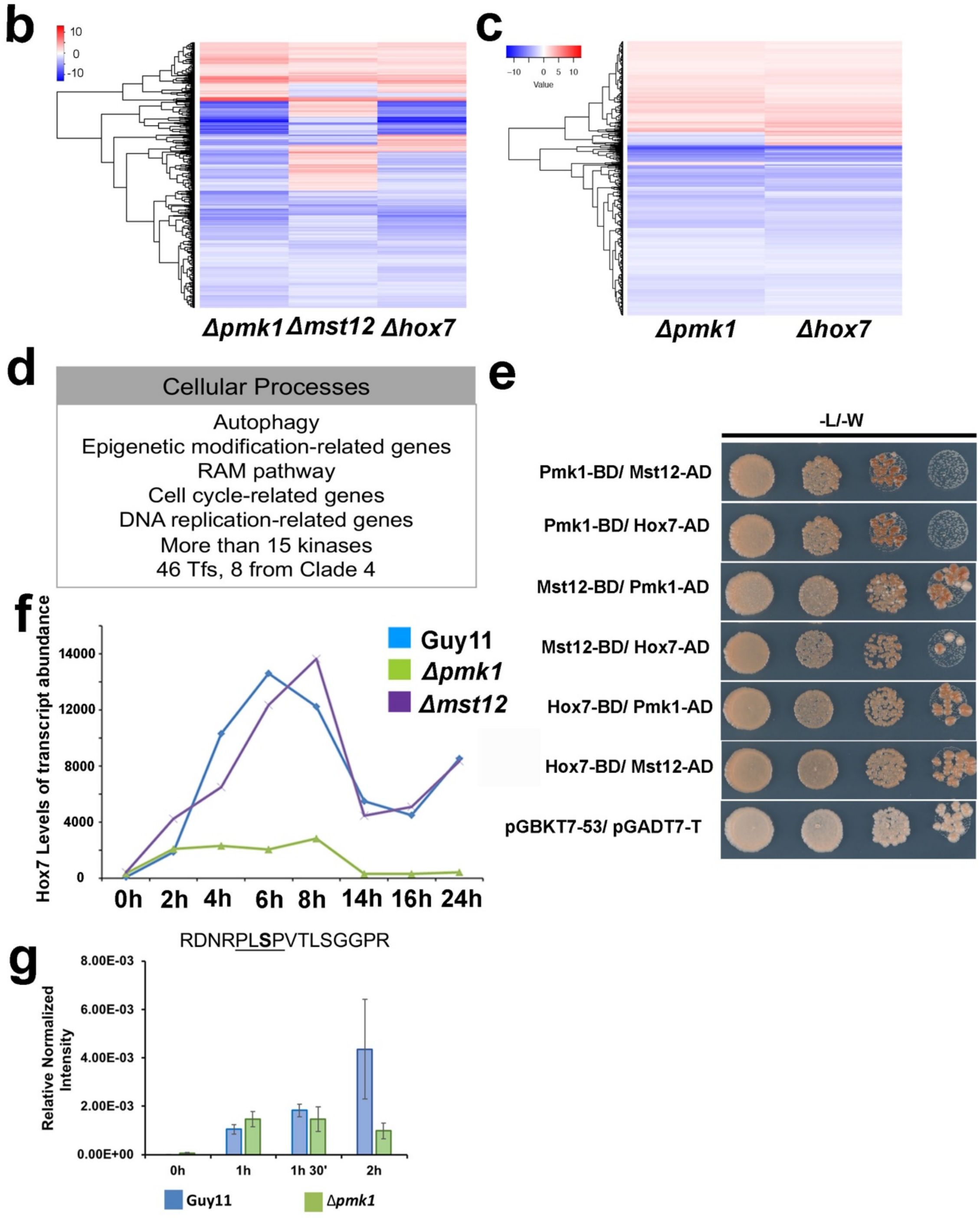


**Supplementary figure 12: Hox7 operates downstream of Pmk1 MAP kinase to regulate a subset of cellular pathways required for appressorium development and plant infection.** a. Table to show the total number of differentially regulated genes, up-regulated genes (padj <0.01; mod-lfc >1) and down-regulated genes (padj <0.01; mod-lfc <-1) of Δ*pmk1 vs.* Guy11, Δ*mst12 vs.* Guy11 and Δ*hox7 vs.* Guy11 at 14h during appressorium development. b. Heatmap showing levels of relative transcript abundance between Δ*pmk1 vs.* Guy11, Δ*mst12 vs.* Guy11 and Δ*hox7 vs.* Guy11 of 709 common genes. c. Heatmap showing levels of relative transcript abundance between Δ*pmk1 vs.* Guy11 and Δ*hox7 vs.* Guy11 of 1942 common genes and most representative cellular pathways found in this gene set. d. Table to show the most representative cellular pathways found in 1942 common genes. e. Yeast-two hybrid analysis of Pmk1, Mst12 and Hox7. Control panel to show the yeast co-transformed an plated in double dropout media. (3 biological replicates). f. Hox7 levels of transcript abundance in Guy11, *Δpmk1* and *Δmst12* during appressorium development time course. g. Parallel Reaction Monitoring (PRM) showing Pmk1-depemdent phosphorylation at serine 158 of the miss-cleaved Hox7 peptide during appressorium development up to 2h after conidial germination.
