## Supplementary Table 9 for "A hierarchical transcriptional network controls appressorium-mediated plant infection by the rice blast fungus *Magnaporthe oryzae*"

**Supplementary Table 4: ­ Primers used in this study**

| Name | Sequence |
| --- | --- |
| Rpp1_50.1 | AGCCGTCCTTGCCGTGGTCG |
| Rpp1_M13F | GTCGTGACTGGGAAAACCCTGGCGAGAAAATCAAGACGCTCCCGCAAT |
| Rpp1_M13R | TCCTGTGTGAAATTGTTATCCGCTATAGTCTCACCGTTGATCTGTCAC |
| Rpp1_30.1 | GGATTCATTGCCAACACTTCTCTC |
| Rpp2_50.1 | GAAAAAGAAACCCGCACAAGACAG |
| Rpp2_M13F | GTCGTGACTGGGAAAACCCTGGCGTGTTGCTGCTGAATGTGACTTGAC |
| Rpp2_M13 R | TCCTGTGTGAAATTGTTATCCGCTGGCATGGGGAACATGTATTACTTG |
| Rpp2_30.1 | CAGTAGAGTAGAATATAGCAGCAG |
| Rpp3_50.1 | CGGAATTCTCCTCGT CTCGCGAGCAAGT |
| Rpp3_HygF | GAATAGAGTAGATGCCGACCCACACGGCTGCAATGCTCAT |
| Rpp3_HygR | TTGACCTCCACTAGCTCCAGCCAAGCCGTTGGTGATGTCTTGGCAGG |
| Rpp3_30.1 | AGCTCTAGACAGCGTCCCTTTCACAACCAG |
| Rpp4_50.1 | TGGTGCCTGTTCCTTGCCTAGC |
| Rpp4_M13F | GTCGTGACTGGGAAAACCCTGGCGACTGCGGTGAGGATTGACGGTG |
| Rpp4_M13R | TCCTGTGTGAAATTGTTATCCGCTTGATTGAGAACCAGCAGGCGTG |
| Rpp4_30.1 | CGGACGTTGGTCTCGGCATTCA |
| Rpp5_50.1 | TTGGTGGCAGTCGCGGAAGAAG |
| Rpp5_M13F | GTCGTGACTGGGAAAACCCTGGCGCGTACCCCTTGACCTACCTTGG |
| Rpp5_M13R | TCCTGTGTGAAATTGTTATCCGCTGGAGGGCTCTGTGATGACTGAT |
| Rpp5_30.1 | ATGTTCGGGGCACCGTGGTTTT |
| Pig1_M13Rv2 | TCCTGTGTGAAATTGTTATCCGCTCGGGATCAAAAGAAAGCTCGCC |
| Pig1_50.1 | GCTCCTCAATCCCACTCCGTCT |
| Pig1_m13F | GTCGTGACTGGGAAAACCCTGGCGTCGGGATGCCAAAAAAGAGGAAAC |
| Pig1_m13R | TCCTGTGTGAAATTGTTATCCGCTTTCCAATTCTTCCACATTAGGCTG |
| Hox7_50.1 | TTTTCTTTGGTCAGTCATTCCGCA |
| Hox7_30.1 | TCATTCCCTATCACGTTTTTGGAG |
| Hox7_M13F | GTCGTGACTGGGAAAACCCTGGCGTGTGGACATTGTGGGTGAATTGCT |
| Hox7_M13R | TCCTGTGTGAAATTGTTATCCGCTTTTATCACGATGACATGCACAGAC |
| Pmk1_F_bait | CATGGAGGCCGAATTCATGTCTCGCGCCAATCCACCAAGC |
| Pmk1_R_bait | GCAGGTCGACGGATCCTTACCGCATAATTTCCTGGTAGAT |
| Mst12_f_prey | GGAGGCCAGTGAATTCATGTATTCCCATCCACACAACGCC |
| Mst12_r_prey | CGATGCCCACCCGGGTCACATCATGCCACCGGCATTGAT |
| Mst12_f_ bait | CATGGAGGCCGAATTCATGTATTCCCATCCACACAACGCC |
| Mst12_r_ bait | GCAGGTCGACGGATCCTCACATCATGCCACCGGCATTGAT |
| Hox7_f_prey | GGAGGCCAGTGAATTCATGGAATATACCTTGCCAAACCAC |
| Hox7_r_prey | CGATGCCCACCCGGGTTAGACGCTGCCACGCTTCATGCC |
| Pmk1_F_prey | GGAGGCCAGTGAATTCATGTCTCGCGCCAATCCACCAAGC |
| Pmk1_R_prey | CGATGCCCACCCGGGTTACCGCATAATTTCCTGGTAGAT |
| Hox7_f_bait | CATGGAGGCCGAATTCATGGAATATACCTTGCCAAACCAC |
| Hox7_r_bait | GCAGGTCGACGGATCCTAGACGCTGCCACGCTTC ATGCC |
| BASplit | GGACTTCAGCAGGTGGGTGTAGAG |
| ARSplit | GCAGACAGGAACGAGGACATTA |
| HYSplit | GGATGCCTCCGCTCGAAGTA |
| YGSplit | CGTTGCAAGACCTGCCTGAA |
| Bar_flp1 | GAGATTTAGGTCGACCAAGAACAACTACCGCTTTCG |
| Flp1_gfp | GCCCTTGCTCACCATCAGACCCGTAATAATAGCGA |
| Bar_flp2 | GAGATTTAGGTCGACCACATATACTTTGCAGCCTGCC |
| Gfp_flp2 | GCCCTTGCTCACCATCAGCTCACCAATGTCCTCCTCG |
