## Supplementary Table 10 for "A hierarchical transcriptional network controls appressorium-mediated plant infection by the rice blast fungus *Magnaporthe oryzae*"

**Supplementary Table 10: Peptides used in PRM assay**

|  | ***Precursor m/z*** | ***Precursor Charge (z)*** |
| --- | --- | --- |
| **HOX7 peptide** |  |  |
| DNRPLS[+80]PVTLSGGPR | 549.2734 | 3 |
| **Control peptides** |  |  |
| VDS[+80]AVQGLSSSPKDTK | 566.9364 | 3 |
| VHS[+80]PVYGGPPGAADSR | 549.5823 | 3 |
| KPAGS[+80]PAVVDKR | 435.5623 | 3 |
| ANVGSM[+16]ADAGDEAM[+16]DVDT[+80]DDDDSNTKKR | 764.5499 | 4 |
| AFGSGPM[+16]S[+80]DEDDEDGHPHGAER | 803.6289 | 3 |
| RRS[+80]PGDGEGDGSVDNR | 585.2465 | 3 |
| IAQAVGGGS[+80]DDDLQAR | 826.8674 | 2 |
| SGS[+80]PQTGTLRQDATTPTLATSVGPSM[+16]DR | 976.7850 | 3 |
| T[+80]KSESKVPAVEPAESR | 598.9593 | 3 |
| ANS[+80]PPPSSLLNHNK | 519.2470 | 3 |
| RKS[+80]SSGSSYAGSYQGTR | 620.2742 | 3 |
| SIS[+80]PGLHAQR | 573.2768 | 2 |
| KVEDEDDGEIS[+80]EPEDPM[+16]M[+16]FQR | 870.0083 | 3 |
| ALDNAPS[+80]GRVT[+80]PVAPPPR | 658.9781 | 3 |
| GM[+16]APYPNS[+80]PQPR | 705.7972 | 2 |
| TSTS[+80]GTATPTAAGR | 679.8010 | 2 |
| SAGREDAQGVIHIDDS[+80]DDEGDVQM[+16]GGTPGPR | 820.0979 | 4 |
| RGSLST[+80]NGSESIDESAIDEDEVGAPNSR | 991.4299 | 3 |
| S[+80]GSITENIFESER | 774.8325 | 2 |
| SDS[+80]HQAGGGVSGDATSLPTLNPVLSR | 868.4081 | 3 |
| NYIFGDPDS[+80]EDEVPR | 916.8724 | 2 |
