## Supplementary Table 6 for "A hierarchical transcriptional network controls appressorium-mediated plant infection by the rice blast fungus *Magnaporthe oryzae*"

| **Name** | **Accession Number** | **Reported Phenotype** | **Reference** |
| --- | --- | --- | --- |
| *ACLR* | MGG_02129 | - | This study |
| *RPP1* | MGG_10212 | - | This study |
| *FZC75* | MGG_05845 | Zn_2_Cys_6_ | Lu *et al.,* 2014 |
| *MoAP1* | MGG_12560 | Oxidative stress response | Guo *et al.,* 2011 |
| *FZC64* | MGG_00320 | Stress response affected | Lu *et al.,* 2014 |
| *FZC50* | MGG_09273 | Conidial germination reduced | Lu *et al.,* 2014 |
| *RPP5* | MGG_08917 | - | This study |
| *RPP4* | MGG_07368 | - | This study |
| *FZC52* | MGG_09676 | Stress response affected | Lu *et al.,* 2014 |
| *PIG1* | MGG_07215 | Putative melanin regulator | Sweigard *et al.,* 1988 |
| *RPP2* | MGG_09276 | - | This study |
| *HOX7* | MGG_12865 | Required for appressorium formation | Kim *et al.,* 2009 |
| *FZC41* | MGG_07534 | Stress response affected | Lu *et al.,* 2014 |
| *FZC30* | MGG_01887 | Stress response affected | Lu *et al.,* 2014 |
| *KV50* | MGG_07218 | - | B. Valent (unpublished) |

**Supplementary Table 3: ­ Table showing the gene name, accession number, reported phenotype and reference of each transcription factors belonging to clade 4.**
